# Discrete genetic effects of *VHL* and *PBRM1* inactivation co-operate to disrupt epithelial homeostasis and promote ccRCC

**DOI:** 10.64898/2026.02.18.706657

**Authors:** Samvid Kurlekar, Joanna D.C.C. Lima, Niklas Kupfer, Christopher W. Pugh, David R. Mole, Julie Adam, Peter J. Ratcliffe

## Abstract

Inactivation of *VHL* is a truncal alteration in clear cell renal cell carcinoma, but additional events are required for oncogenesis, most commonly *PBRM1* inactivation. To better understand this co-operation, we used an oncogenic cell-tagging strategy to analyze the earliest transcriptional and cellular consequences of *Vhl* and/or *Pbrm1* inactivation in the renal tubular epithelium, *in vivo,* at single-cell resolution. *Pbrm1* inactivation did not globally alter HIF-dependent transcription or increase early tubular proliferation induced by *Vhl* inactivation. Instead, it had independent effects on epithelial organization. Combined genetic and morphological analyses suggested that *Pbrm1* inactivation allows cells to sustain *Vhl*/HIF-dependent proliferation by disrupting tubular architectures that ordinarily restrain this proliferation, resulting in extra-tubular cell accumulation, multilayered epithelia, and tumor formation. Our findings frame a new model for the *VHL*-*PBRM1* interaction that explains loss of epithelial homeostasis through an interaction between discrete effects that drive proliferation and remove structural tissue restraints on that proliferation.

**Statement of Significance:** *VHL* and *PBRM1* are frequently co-inactivated in ccRCC. Combining transcriptomic and histological analyses of *Vhl* and/or *Pbrm1*-inactivated renal cells in vivo, this study highlights independent effects on transcription and epithelial organization that converge to promote sustained proliferation and dysplasia. The work illuminates how tissue homeostasis is disrupted by oncogenic co-operation.

## Introduction

Large-scale sequencing of tumor DNA has identified mutations in numerous genes that provide robust entry points to understanding cancer. A cardinal finding of this work is the interplay between mutations in different genes, which is poorly understood. The high prevalence and specificity of many such interactions suggests that they should provide insights of general importance to cancer. In this work, we took as an example two commonly observed sites of mutation in clear cell renal cell carcinoma (ccRCC); the gene encoding the VHL (von Hippel Lindau) ubiquitin E3 ligase and the gene encoding PBRM1 (Polybromo-1), the nucleosome binding subunit of the PBAF SWI/SNF chromatin remodeling complex (1). Both genes behave as tumor suppressors in this context (1).

Biallelic inactivating mutations in *VHL* are observed in at least 80% of ccRCC, whereas such mutations in *PBRM1* are observed in approximately 40% of ccRCC, generally occurring after *VHL* inactivation (2,3). Though other functions have been proposed for VHL, its best-defined activity is as an E3 ubiquitin ligase targeting Hypoxia Inducible Factors (HIFs) (4,5). HIFs are transcription factors that transduce an extensive transcriptional response to hypoxia. In VHL-defective cells, the oxygen-dependent degradation of HIFα subunits is disabled, leading to constitutive HIF activation regardless of oxygen levels (4).

In spontaneous cases of human ccRCC, the initiating event is most commonly loss or re-organization of chromosome 3p that disrupts both *VHL* and *PBRM1*, followed by inactivation of the second *VHL* allele (6). Patients with VHL disease, who exhibit increased susceptibility to a range of tumors including ccRCC, inherit one defective copy of *VHL* while the other is lost somatically, generally via 3p loss (6). In affected tumors of both types, loss of the second *PBRM1* allele occurs subsequently, by mutational inactivation (6). In mice, *Vhl* and *Pbrm1* lie on different chromosomes so cannot be co-deleted by a single event and *Vhl* inactivation alone does not cause tumors (7). However, simultaneous inactivation of *Vhl* and *Pbrm1* by expression of *Cre recombinase* restricted to the renal tubular epithelium (RTE) results in multiple tumors within the renal cortex, with histological features resembling those of human ccRCC (8–10). As with the human disease, these studies reveal substantial latency in the development of RCC. Together with the high prevalence of mutations in both *VHL* and *PBRM1* in human RCC, this work emphasizes the causal importance of co-inactivation of *VHL* and *PBRM1* in the development and progression of RCC and has generated great interest in understanding this interaction.

PBRM1 and the PBAF complex have been implicated in regulating transcription (11), DNA repair (12), and the resolution of replicative stress (8). Non-nuclear functions in microtubule organization (13) and epithelial integrity (14,15) have also been described. However, given the role of VHL in regulation of gene expression via the HIF pathway and that of PBRM1 in chromatin remodeling, many studies of its oncogenic role in RCC have focused on the possibility that PBRM1 directly affects patterns of gene expression driven by *VHL* inactivation, such as the HIF transcriptional cascade. These studies have arrived at different conclusions; for instance, enhancement (10,16), inhibition (14), isoform-specific modulation (17) and no effect (3) on the HIF transcription response, have all been reported. Thus, despite their frequent co-inactivation in ccRCC and intense investigation, the interaction between *VHL* and *PBRM1* inactivation in oncogenesis remains poorly understood.

Development of an oncogenic cell tagging mouse model of *Vhl* inactivation has enabled the in vivo cellular and transcriptomic consequences of *Vhl* inactivation to be resolved in the RTE with spatiotemporal resolution (18). In this model, the conditional excision of an engineered *Vhl* allele is directly linked to the activation of a tdTomato reporter. When this allele is coupled with a constitutively inactive *Vhl* allele, tamoxifen administration to promote recombination in early adult life mimics the ‘second hit’ in familial VHL disease (18,19). Using this model, we reported previously that changes in gene expression associated with *Vhl* inactivation were, as expected, dominated by changes directly or indirectly dependent on HIF, but remarkably specific to different tubular cell types (20). We also observed a proliferative phenotype in proximal tubular (PT) cells, the proposed cells-of-origin for ccRCC (21,22), that required both HIF1A and HIF2A isoforms (20). Surprisingly, however, despite sustained activation of HIF, this proliferation was transient and apparently constrained by the structural integrity of the normal renal tubule, with proliferating cells remaining confined within a single epithelial layer bounded by the basement membrane (18).

In the present work, we extended the oncogenic cell labeling strategy to investigate the consequences of combined *Vhl* and *Pbrm1* inactivation by breeding the cell tagging mouse model to carry additionally a conditionally inactivated *Pbrm1* gene enabling biallelic inactivation of both *Vhl* and *Pbrm1* upon tamoxifen administration. This allowed us to directly compare transcriptional and morphological changes in *Vhl*- versus *Vhl/Pbrm1*-inactivated RTE cells in vivo with single-cell resolution. Importantly, because affected *Vhl*-inactivated cells were tagged with tdTomato, altered phenotypes could be studied in advance of overt tumor formation at a time when *Pbrm1* inactivation first exerts tumorigenic effects in *Vhl*-null cells. Our findings revealed that *Pbrm1* inactivation had little effect on the HIF transcriptional response. Rather, we observed largely distinct effects of *Pbrm1* inactivation, including changes in the expression of genes associated with epithelial integrity, cell motility and lipid metabolism. Surprisingly, early proliferation was not increased upon *Pbrm1* inactivation but was strikingly more sustained than with *Vhl* inactivation alone. This was associated with disruption of the renal tubular anatomy, with the appearance of dysplastic lesions manifesting a multi-layered epithelium, with the accrual of cells within the tubular lumen and appearance of cells crossing the basement membrane to occupy an interstitial location.

Overall, our findings suggest that rather than interacting at the level of gene expression, oncogenic inactivations of *VHL* and *PBRM1* interact at the level of the cellular phenotype, with *PBRM1* inactivation appearing to release cells from the anatomical confines of renal epithelial integrity and resulting in maintenance of the proliferation that is induced by *VHL* inactivation, multiple epithelial abnormalities and renal tumors resembling ccRCC.

## Results

### Co-deletion of *Pbrm1* in an oncogenic ‘cell tagging’ model of *Vhl* inactivation

To assay the effects of concomitant *Vhl* and *Pbrm1* inactivation, we bred ‘floxed’ *Pbrm1* alleles (*Pbrm1^fl/fl^*) into a previously reported oncogenic cell tagging mouse model of *Vhl* inactivation (18) (**Supplementary Fig. 1a**). In this model, a *Vhl* allele (*Vhl^pjr.fl^*) that links conditional *Vhl* knock-out directly to tdTomato knock-in is coupled with either a wild-type (*Vhl^wt^*) or constitutively knocked-out (*Vhl^jae.KO^*) (23) untagged second *Vhl* allele. Tamoxifen-induced and RTE-restricted Cre-mediated recombination with *Pax8-CreERT2* (24) generates tdTomato-tagged, *Vhl*-null proximal tubular (PT) cells in the renal cortex and outer medulla, and collecting duct cells in the renal papilla in *Vhl^jae.KO/pjr.fl^; Pax8-CreERT2* (‘VKO’) mice. These were compared to tdTomato-tagged but *Vhl*-haplosufficient cells of matching types in *Vhl^wt/pjr.fl^; Pax8-CreERT2* (‘ConKO’) mice (18,20). By combining this model with floxed *Pbrm1* alleles in *Vhl^jae.KO/pjr.fl^; Pbrm1^fl/fl^; Pax8-CreERT2* (‘VPKO’) and *Vhl^wt/pjr.fl^; Pbrm1^fl/fl^; Pax8-CreERT2* (‘ConPKO’) mice, we sought to generate tdTomato-tagged *Vhl*-null and *Vhl*-haplosufficient RTE cells respectively that were also *Pbrm1*-null (**Table 1** and **Supplementary Fig. 1a**).

**Table 1:** Genotype nomenclature, and *Vhl* and *Pbrm1* status in tdTomato-positive and tdTomato-negative cells in mice used in this study.

| Abbreviation | Genotype | tdTomato-negative Cells |  | tdTomato-positive Cells |  |
| --- | --- | --- | --- | --- | --- |
|  |  | *Vhl* | *Pbrm1* | *Vhl* | *Pbrm1* |
| ConKO | *Vhl*<sup>wt/pjr.fl.</sup>,<br>*Pbrm1*<sup>wt/wt.</sup>,<br>*Pax8*-<br>*CreERT2* | Both alleles functional | Both alleles functional | One allele inactivated (tagged) | Both alleles functional |
| ConPKO | *Vhl*<sup>wt/pjr.fl.</sup>,<br>*Pbrm1*<sup>fl/fl.</sup>,<br>*Pax8*-<br>*CreERT2* | Both alleles functional | Both alleles functional | One allele inactivated (tagged) | Both alleles inactivated |
| VKO | *Vhl*<sup>jae.KO/pjr.fl.</sup>,<br>*Pbrm1*<sup>wt/wt.</sup>,<br>*Pax8*-<br>*CreERT2* | One allele inactivated (untagged) | Both alleles functional | Both alleles inactivated (tagged) | Both alleles functional |
| VPKO | *Vhl*<sup>jae.KO/pjr.fl.</sup>,<br>*Pbrm1*<sup>fl/fl.</sup>,<br>*Pax8*-<br>*CreERT2* | One allele inactivated (untagged) | Both alleles functional | Both alleles inactivated (tagged) | Both alleles inactivated |

We tested whether the floxed *Pbrm1* alleles recombined fully and concomitantly with the cell tagging *Vhl* allele by assaying the genomic status and protein levels of PBRM1 in flow-sorted tdTomato-positive and negative renal cells (see sorting strategy in **Supplementary Fig. 1b**, biased towards the isolation of PT cells (18,20)). In tdTomato-positive cells from VPKO mice given 5x 2 mg TMX and harvested 1-3 weeks following recombination (‘early’ timepoint), the unrecombined form of the *Pbrm1* allele was not detected by genomic PCR (**Supplementary Fig. 1c**) and PBRM1 protein was absent in immunoblotting (**Supplementary Fig. 1d**). In contrast, the wild-type or unrecombined forms of the *Pbrm1* allele were present, and PBRM1 protein was detected in tdTomato-negative cells from VPKO mice and all cells from VKO mice respectively. These data indicated that tdTomato-positivity, and by extension, *Vhl^pjr.fl^* recombination was accurately coupled to *Pbrm1^fl^* recombination in our model.

We then tested if concomitant *Vhl* and *Pbrm1* inactivation in this mouse model led to the formation of renal tumors. Multiple tumors were observed in the renal cortex of VPKO mice (n = 4/4) harvested 12-17 months following tamoxifen recombination. They exhibited cell clearing and stained positively for tdTomato (**Supplementary Fig. 1e**), were positive for HIF1A and HIF2A, and did not express PBRM1 (**Supplementary Fig. 1f**). These tumors were morphologically similar to, though somewhat smaller than, the renal tumors reported by others to have formed 20-24 months following concomitant *Vhl* and *Pbrm1* inactivation in the PT (8). Taken together, these results indicated that this mouse model could generate renal tumors, that the tumorigenic effects of *Vhl* and *Pbrm1* inactivation were manifest by 12 months following recombination, and that tdTomato-positivity accurately reported concomitant *Vhl* and *Pbrm1* loss. Having established this, we focused on defining the earliest contributions of *Vhl* and/or *Pbrm1* inactivation to gene expression and epithelial phenotypes in PT cells in vivo to gain mechanistic clues as to the pathogenesis of renal cancer.

### Effect of *Pbrm1* inactivation on *Vhl* and HIF-dependent gene expression in PT cells

*PBRM1* inactivation has been linked previously to both amplification and diminution of HIF-dependent gene expression in ccRCC cell lines and in kidneys from mouse models in which *Vhl* and *Pbrm1* are inactivated in renal tubules in utero (10,14,16). We sought to address this controversy by testing for a positive or negative effect on the HIF response. To do so, we performed single-cell RNA sequencing (scRNA-seq) on tdTomato-positive cells from kidneys of VPKO mice harvested 1-3 weeks following recombination (early timepoint). This data was integrated with that obtained previously from tdTomato-positive cells from kidneys of ConKO and VKO mice harvested at a similar timepoint (18,20). After integration, the dataset contained 149,550 high-quality cells from mice of both sexes and at least n = 3 mice per genotype, with an average of 13,595 ± 7,346 (SD) cells per mouse, 5,127 ± 2,065 (SD) mapped reads per cell, and 1,690 ± 368 (SD) genes detected per cell.

Uniform manifold approximation and projection (UMAP) and Louvain clustering analysis revealed that ConKO and VKO cells occupied largely non-overlapping UMAP space and clusters, consistent with the known substantial effects of *Vhl* inactivation on the transcriptome (**Fig. 1a, b**). Notably, in the same analysis, VPKO cells occupied distinct UMAP space and clusters from VKO and ConKO cells, indicating that *Pbrm1* inactivation had induced further transcriptomic changes in these cells (**Fig. 1a, b**).

**Fig. 1:**
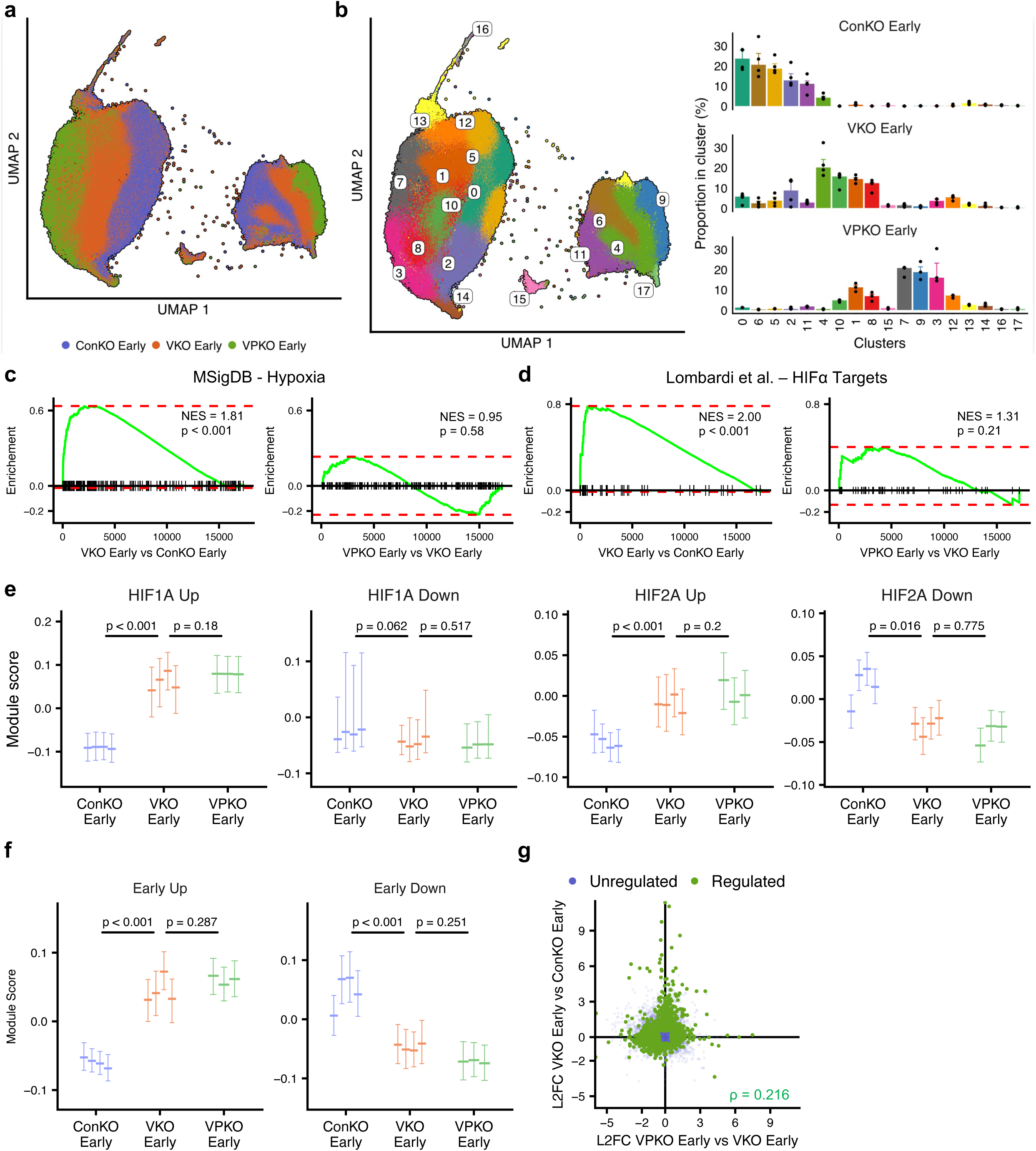
Effect of *Pbrm1* inactivation on *Vhl* and HIFα-dependent gene expression in PT cells. **(a)** UMAP plot depicting ConKO, VKO, and VPKO cells harvested early (1-3 weeks) after recombination, showing that VPKO cells occupy distinct UMAP space to VKO and ConKO cells. **(b)** Cell clusters occupied by ConKO, VKO, and VPKO cells in UMAP space (left panel) and the proportion of cells from mice of each genotype that are present in each cluster (right panel). Data are presented as median values, with the inter-quartile range indicated by error bars. **(c, d)** Gene-set enrichment analyses of the MSigDB Hallmark Hypoxia pathway **(c)** or HIFα target genes as defined in Lombardi et al., 2022 **(d)** among differentially regulated genes in VKO Early versus ConKO Early cells (left panel) or VPKO Early versus VKO Early cells (right panel). NES – normalized enrichment score. P value adjusted by the false discovery rate. **(e)** Expression of sets of genes up- or downregulated in cells of any PT identity in a HIF1A-(‘HIF1A Up’ or ‘HIF1A Down’) or HIF2A-dependent manner (‘HIF2A Up’ or ‘HIF2A Down’) in ConKO, VKO, and VPKO cells sequenced early (1-3 weeks) following recombination. **(f)** Expression of sets of genes up- or downregulated in cells of any PT identity early after *Vhl* inactivation (‘Early Up’ or ‘Early Down’) in ConKO, VKO, and VPKO cells sequenced 1-3 weeks (early) following recombination. In **(e, f**) median values and inter-quartile ranges of the data are presented separately for cells from each mouse. Pairwise comparisons have been made between means of the median values for mice of different genotypes using two-tailed one-way ANOVA with multiple testing correction using the Benjamini-Hochberg method. **(g)** Pseudo-bulked log_2_-fold changes (L2FC) in VPKO Early versus VKO Early cells plotted against changes in VKO Early versus ConKO Early cells. Spearman’s correlation coefficient (ρ) calculated for genes that exhibit statistically significant regulation in either of the depicted comparisons.

To assess the contribution of additional *Pbrm1* inactivation to HIF-dependent gene expression, we performed pseudo-bulked differential expression analysis in VPKO versus VKO cells or in VKO versus ConKO cells (data provided as a resource in **Supplementary Table 1**) and analyzed for the enrichment of either a publicly curated hypoxia pathway gene set, or a published set of pan-cancer direct HIF target genes (25). As expected, both gene sets were significantly upregulated in VKO versus ConKO cells, but neither was significantly enriched in VPKO versus VKO cells (**Fig. 1c, d**). Therefore, in our analyses, *Pbrm1* inactivation neither amplified nor diminished the collective expression of hypoxia- or HIF-dependent genes as have been defined in cancer cells.

In previous work, we had identified sets of established and novel genes that were regulated in vivo in PT cells in a HIF1A- or HIF2A-dependent manner (20). To test for an effect on HIF-dependent genes in this setting, we scored ConKO, VKO, and VPKO cells at an early timepoint (1-3 weeks following recombination) for the expression of these genes (20). While the expression of each of the HIF1A- and HIF2A-dependent up- or downregulated gene sets was significantly altered in VKO versus ConKO cells as expected, none of these HIF-dependent gene sets was altered further in VPKO versus VKO cells (**Fig. 1e**). PT cells can be divided into three ‘types’ – S1, S2, and S3 – that differ in physiological function, anatomical location, and gene expression (26). PT cells of each type have been further categorized into two classes – A and B – the former of which is characterized by specific expression of AP-1 transcription factors and long non-coding RNAs such as *Neat1* and *Malat1* (18). Together, PT type and class define six PT ‘identities’ – S1 A, S1 B, S2 A, S2 B, S3 A and S3 B – that differ in their isoform-specific HIF-dependent response to *Vhl* inactivation (20). Therefore, we also tested if HIF-dependent genes were altered specifically in any PT identity. When cells were analyzed separately for the expression of HIF1A- and HIF2A-dependent genes specific to their PT identity, an effect of *Vhl* inactivation was evident as expected, but no significant differences were observed between VPKO versus VKO cells (**Supplementary Fig. 2a**). Therefore, we found no evidence that *Pbrm1* inactivation impacted the expression of HIF-dependent genes either overall or within any specific subtype of PT cells in vivo.

To test if *Pbrm1* inactivation altered the transcriptional response to *Vhl* inactivation in any way, irrespective of HIF-dependence, we scored cells for the expression of genes previously shown (18) to be up- and downregulated early following *Vhl* inactivation. While both gene sets were significantly altered in VKO versus ConKO cells, no further alterations were observed in VPKO versus VKO cells (**Fig. 1f**). No differences were observed even when cells were analyzed separately for the expression of early *Vhl*-dependent genes specific to their PT identity (**Supplementary Fig. 2b**). Consistent with this, pseudo-bulked differential gene expression in VPKO versus VKO cells exhibited poor correlation to that in VKO versus ConKO cells (**Fig. 1g**).

These results indicated that the early effects of *Pbrm1* inactivation are not simple enhancements or diminutions of the HIF transcriptional response and are largely distinct from the effects of *Vhl* inactivation alone.

### Sustained and independent transcriptional effects of *Pbrm1* inactivation in PT cells

To understand the biological roles of genes affected by *Pbrm1* inactivation, we assessed the over-representation of Gene Ontology (GO) terms among genes significantly regulated (|log_2_-fold change (L2FC)| > 0.5, p < 0.01) in pseudo-bulked differential expression analysis of VPKO versus VKO cells harvested early (1-3 weeks) following recombination. To gauge the magnitude of these effects, we evaluated the average L2FC across all genes comprising each over-represented GO term. Among upregulated genes, several genes and GO terms pertaining to lipid and sterol metabolism (e.g., *Pcsk9*, *Srebf1*, *Insig1*) were significantly (p < 0.01, average L2FC > 0) over-represented (**Fig. 2a, b**). Interestingly, several genes and terms relating to epithelial and tissue morphogenesis, cell adhesion, and motility (e.g., *Cldn1*, *Itga9*, *Adamts15*, *Nid2*) were also over-represented (**Fig. 2a, b**). No GO terms were significantly over-represented within downregulated genes, although several downregulated genes had known roles in epithelial cell polarity, adhesion, and morphogenesis, such as *Cdc42ep5* (27)*, Trp53bp2* (28), and *Cldn34c1* (29) (**Fig. 2a**).

**Fig. 2:**
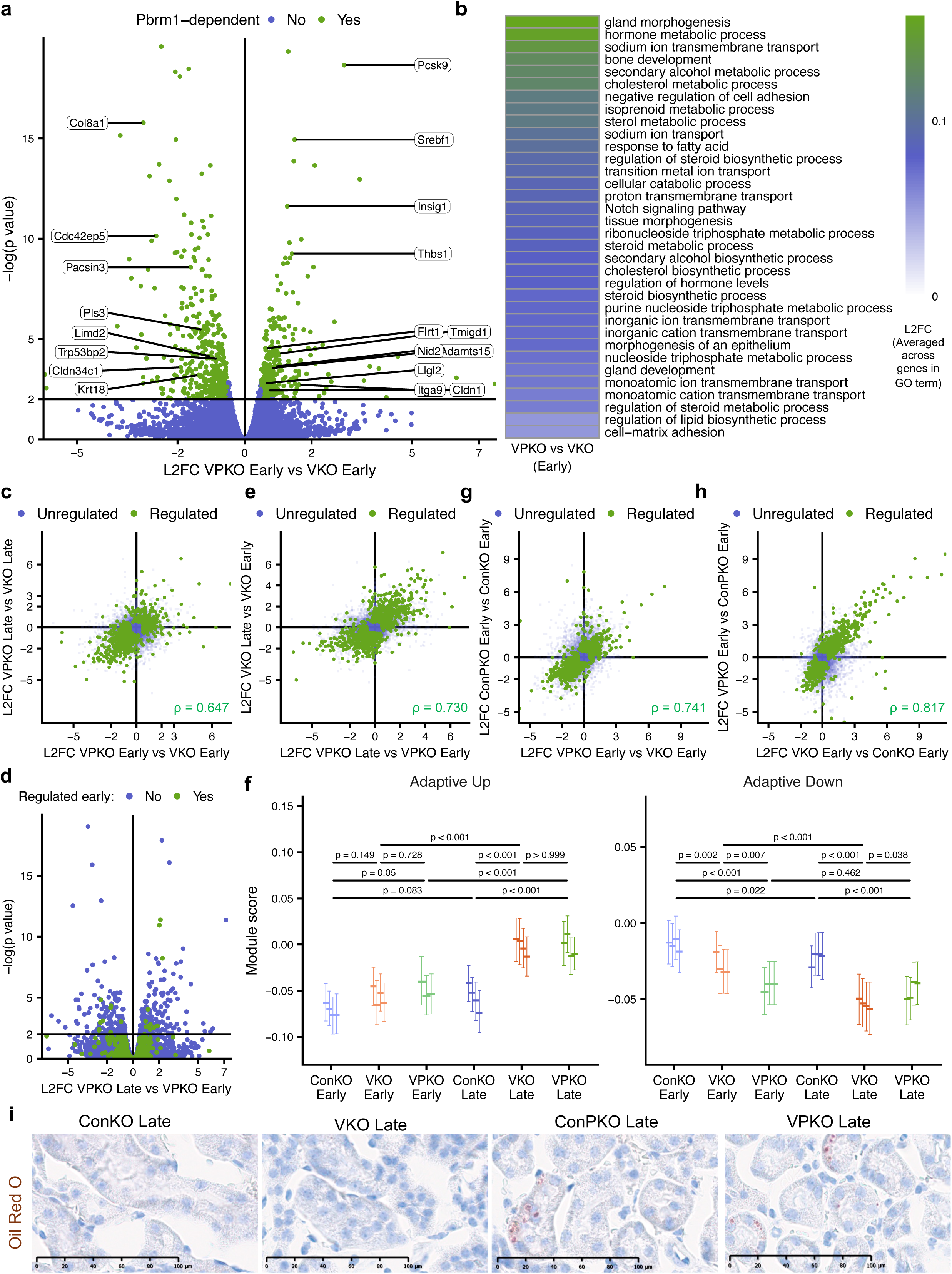
Sustained and independent transcriptional effects of *Pbrm1* inactivation in PT cells. **(a)** Pseudo-bulked L2FC plotted against the negative logarithm of the p value (two-tailed Wald test with multiple testing correction by Benjamini-Hochberg method) for genes in VPKO Early versus VKO Early cells. Genes significantly (p < 0.01) and robustly (|L2FC| > 0.5) regulated are colored green. Candidate genes pertaining to lipid and sterol metabolism and epithelial morphogenesis and adhesion are labelled. **(b)** Gene Ontology terms for biological processes that are significantly over-represented among genes upregulated by *Pbrm1* inactivation in *Vhl*-null cells. GO terms whose over-representation is significant (p < 0.01; one-sided Fisher’s exact test corrected by false discovery rate), and whose member genes together exhibit a net positive average L2FC in VPKO Early versus VKO Early cells are shown. Terms are ordered and tiles are colored by the average L2FC. **(c)** Pseudo-bulked L2FC in VPKO Early versus VKO Early cells plotted against changes in VPKO Late versus VKO Late cells showing that *Pbrm1*-dependent regulation of genes is correlated between the two timepoints. **(d)** Pseudo-bulked L2FC plotted against the negative logarithm of the p value (two-tailed Wald test with multiple testing correction by Benjamini-Hochberg method) for genes in VPKO Late versus VPKO Early cells, showing that genes significantly regulated after *Pbrm1* inactivation at the early timepoint (colored green) are mostly not regulated significantly over time (i.e., have either |L2FC| < 0.5 or p > 0.01 in the VPKO Late versus VPKO Early comparison). **(e)** Pseudo-bulked L2FC in VPKO Late versus VPKO Early cells plotted against changes in VKO Late versus VKO Early cells showing that time-dependent changes in gene expression are correlated in *Vhl-*null and *Vhl/Pbrm1*-null cells. **(f)** Expression of sets of genes up- or downregulated over time in *Vhl*-null cells of any PT identity (‘Adaptive Up’ or ‘Adaptive Down’) in ConKO, VKO, and VPKO cells sequenced either 1-3 weeks (early) or 4-12 months (late) following recombination. Median values and inter-quartile ranges of the data are presented separately for cells from each mouse. Pairwise comparisons have been made between means of the median values for mice of different genotypes using two-tailed two-way ANOVA with multiple testing correction using the Benjamini-Hochberg method. Adaptive upregulation is maintained, and adaptive downregulation is partially accelerated following *Pbrm1* inactivation in *Vhl*-null cells. **(g)** Pseudo-bulked L2FC in VPKO Early versus VKO Early cells plotted against changes in ConPKO Early versus ConKO Early cells, showing that *Pbrm1*-dependent gene regulation is correlated in *Vhl*-null and *Vhl*-competent cells. **(h)** Pseudo-bulked L2FC in VKO Early versus ConKO Early cells plotted against changes in VPKO Early versus ConPKO Early cells, showing that *Vhl*-dependent gene regulation is correlated between *Pbrm1*-competent and *Pbrm1*-null cells. **(c, e, g, h**) Spearman’s correlation coefficient (ρ) calculated for genes that exhibit significant regulation in either of the depicted comparisons. **(i)** Oil Red O staining in renal sections from ConKO, ConPKO, VKO, and VPKO mice harvested 4-12 months (late) following recombination, showing the accumulation of lipid droplets (stained red) in PT cells of ConPKO and VPKO mice. Scale bar, 100 μm. 40x magnification.

To assess if changes induced by *Pbrm1* inactivation early after recombination were sustained, we performed scRNA-seq on tdTomato-positive cells from kidneys of VPKO mice harvested between 4-12 months following recombination (‘late’ timepoint) and integrated the data with that obtained previously from tdTomato-positive cells from ConKO and VKO mice at a similar late timepoint (18). After integration, the dataset contained 178,828 high-quality cells from n = 2 male and n = 2 female mice per genotype, with an average of 14,902 ± 6,587 (SD) cells per mouse, 4,282 ± 1,192 (SD) mapped reads per cell, and 1,455 ± 291 (SD) genes detected per cell. After integration with the dataset of cells harvested ‘early’ after recombination, the combined dataset contained 328,378 high-quality cells and enabled analysis of time-dependent changes in *Vhl*-null and *Vhl/Pbrm1*-null cells.

Pseudo-bulked differential gene expression in VPKO Late versus VKO Late cells exhibited strong correlation with that in VPKO Early versus VKO Early cells (**Fig. 2c**). Only a few genes regulated early in a *Pbrm1*-dependent manner exhibited significant (|L2FC| > 0.5, p < 0.01) time-dependent regulation in *Vhl/Pbrm1*-null cells (**Fig. 2d**). These data indicated that early *Pbrm1*-dependent transcriptional differences were largely maintained over time.

Nevertheless, time-dependent changes in gene expression were observed in *Vhl/Pbrm1*-null cells (**Fig. 2d** and **Supplementary Table 1**). Previously, we had noted that *Vhl* inactivation drives time-dependent changes over one year that are marked by upregulation of genes involved in lipid and amino acid metabolism and downregulation of renal differentiation markers (18). We therefore assessed whether time-dependent changes in VPKO cells over a similar timescale resembled those in VKO cells. Pseudo-bulked differential gene expression in VPKO Late versus VPKO Early cells was highly correlated with that in VKO Late versus VKO Early cells, suggesting that the adaptive response to *Vhl* inactivation is retained following *Pbrm1* inactivation (**Fig. 2e**).

To test this further, we scored all cells for expression of genes previously shown to be either up- or downregulated over time specifically in *Vhl*-null cells (18). VKO and VPKO cells exhibited similar time-dependent increases in the adaptive upregulated gene set and attained similar levels of expression at a late timepoint (**Fig. 2f**). Interestingly, genes classified as adaptively downregulated, and which contribute to a de-differentiated phenotype (20), were expressed at lower levels in VPKO versus VKO cells at 1-3 weeks following recombination but did not decrease further over time (**Fig. 2f**). This suggested that *Pbrm1* loss may partially accelerate the dedifferentiation of *Vhl*-null cells.

As the contributions of *Vhl* and *Pbrm1* inactivation to gene expression appeared to be largely separate, we tested if the response to *Pbrm1* inactivation was dependent at all on concomitant *Vhl* inactivation. We performed scRNA-seq on tdTomato-positive cells from ConPKO mice harvested between 1-3 weeks following recombination (early timepoint). Data from 27,795 high-quality cells (n = 2 male and n = 1 female mice), with an average of 9,265 ± 128 (SD) cells per mouse, 7,580 ± 1,004 (SD) mapped reads per cell, and 2,136 ± 143 (SD) genes detected per cell, were integrated with the data from ConKO, VKO, and VPKO mice at a similar early timepoint. UMAP and Louvain clustering revealed that ConPKO cells occupied largely distinct UMAP space and clusters, consistent with *Pbrm1*-dependent transcriptomic changes (**Supplementary Fig. 3a, b**).

Pseudo-bulked differential gene expression in ConPKO versus ConKO cells correlated strongly to that in VPKO versus VKO cells, suggesting that *Pbrm1* inactivation had similar effects in PT cells with or without *Vhl* inactivation (**Fig. 2g** and **Supplementary Table 1**). Conversely, pseudo-bulked differential gene expression in VKO versus ConKO cells also correlated strongly to that in VPKO versus ConPKO cells, suggesting that *Vhl* inactivation elicited similar effects regardless of *Pbrm1* status (**Fig. 2h** and **Supplementary Table 1**). Consistent with changes in expression of genes involved in lipid and sterol metabolism that were induced by *Pbrm1* inactivation irrespective of *Vhl* status, lipid droplets were observed by Oil Red O staining in PT cells of both ConPKO and VPKO mice, but not of ConKO and VKO mice, in a set of animals examined 10-12 months following recombination (**Fig. 2i**).

Taken together, these results indicated that the transcriptional effects of *Pbrm1* inactivation encompass sustained changes in lipid and sterol homeostasis and epithelial tissue organization, and that they occur independently of the effects of biallelic *Vhl* inactivation in PT cells.

### Emergent cell states among *Vhl/Pbrm1*-null cells

The data so far led us to consider whether changes incurred in *Vhl/Pbrm1*-null cells over time can be explained entirely as a summation of the separate early and time-accrued effects of *Vhl* inactivation and the early and sustained effects of *Pbrm1* inactivation. To test this, we analyzed the correlation between pseudo-bulked differential expression in VPKO Late versus ConKO Late cells (‘observed’ time-accrued effects of *Vhl* and *Pbrm1* inactivation) and the summed changes in VKO Late versus ConKO Late and VPKO Early versus VKO Early cells (‘predicted’ changes based on the individual actions of *Vhl* and *Pbrm1* inactivation). These analyses showed that observed and predicted changes were highly correlated, suggesting that the altered expression of most genes in *Vhl/Pbrm1*-null cells could be explained by additive and separate effects of *Vhl* and *Pbrm1* inactivation (**Fig. 3a**). However, a few genes (termed ‘emergent’ genes) exhibited a significantly (p < 0.01 by two-sided Wald z-test) greater magnitude of change in VPKO Late versus ConKO Late cells than predicted. Upregulated emergent genes included members of the integrated stress response (ISR) (30), such as *Atf5* and *Mthfd2* (**Fig. 3a**). Downregulated emergent genes included renal developmental transcription factors such as *Pax2* (31), cell adhesion molecules such as *Arvcf* (32), cytoskeletal genes such as *Csrp1* (33), and those involved in epithelial planar cell polarity such as *Fzd3* and *Shroom3* (34) (**Fig. 3a**).

**Fig. 3:**
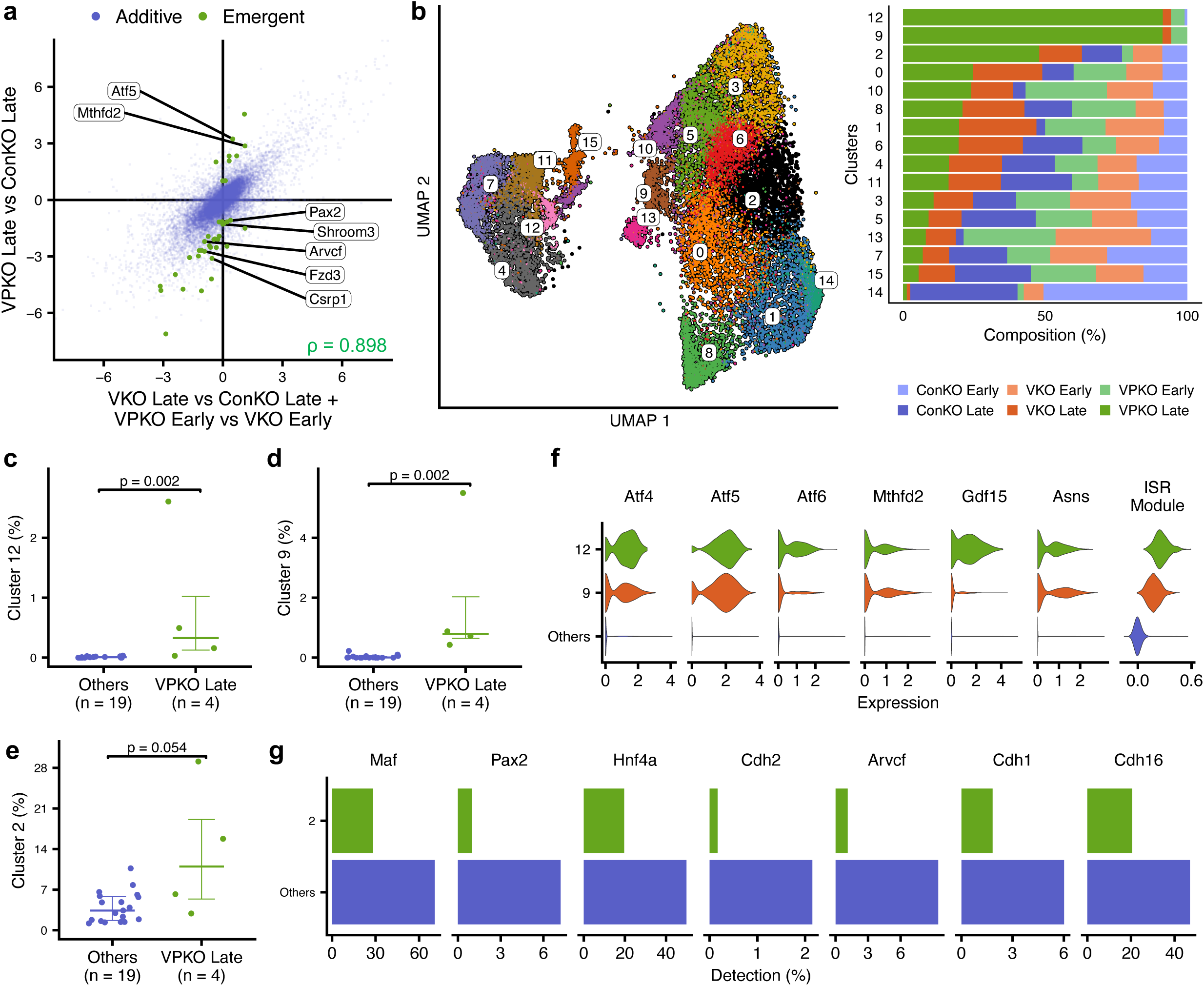
Emergent cell states among *Vhl/Pbrm1*-null cells. **(a)** Pseudo-bulked L2FC in VPKO Late versus ConKO Late cells plotted against ‘predicted’ changes calculated as addition of the effects of *Vhl* inactivation (VKO Late vs ConKO Late) and early effects of *Pbrm1* inactivation (VPKO Early vs VKO Early). Genes that exhibited significant regulation in either of the depicted comparisons are plotted. Genes whose regulation in VPKO Late versus ConKO Late cells exhibit significant (p < 0.01; two-sided Wald z-test) deviation from the additive effects of *Vhl* and *Pbrm1* inactivation are colored green. Spearman’s correlation coefficient (ρ). **(b)** Cells from ConKO, VKO, and VPKO mice projected onto UMAP space and clusters defined for VPKO Late cells (left panel), and the proportion of cells in each cluster derived from mice of each of the genotypes and timepoints (right panel), showing differential occupancy of Clusters 12, 9, and 2 by VPKO Late cells. **(c-e)** Proportion of cells from VPKO Late mice or mice from other genotypes and timepoints in Clusters 12 **(c)**, 9 **(d**), and 2 **(e**). Pairwise comparisons by Wilcoxon test. Error bars indicate the median and the inter-quartile range. **(f)** Expression of candidate genes that are part of the integrated stress response (ISR), and expression score for an ISR gene module derived from Han et al., 2023, in cells from Clusters 9, 12, or other clusters, showing that Clusters 12 and 9 are both characterized by increased expression of ISR genes. **(g)** Proportion of cells with detected reads for candidate genes downregulated in cells from Cluster 2 compared to those in the remaining clusters, showing that Cluster 2 cells are characterized by reduced expression of several PT differentiation transcription factors and adhesion molecules.

Given the paucity of ‘emergent’ genes and their association with specific biological processes, we wondered if they were dysregulated in sub-populations of cells that emerged over time following *Vhl* and *Pbrm1* inactivation. To test this, we characterized cell state heterogeneity among VPKO Late cells by UMAP and Louvain clustering. As expected, VPKO Late cells, all being of the same genotype and timepoint, segregated mainly according to their PT cell type and PT class (**Supplementary Fig. 4a**). To identify clusters disproportionately occupied by VPKO Late cells, we projected cells from other genotypes and timepoints onto the VPKO Late UMAP space and analyzed cluster composition. This identified three clusters – 12, 9, and to a lesser extent 2 – as being disproportionately occupied by VPKO Late cells, suggesting that these represented cell states which emerged in *Vhl/Pbrm1*-null cells over time (**Fig. 3b-e**).

Cluster-wise differential expression analysis revealed that Clusters 12 and 9, composed of PT Class A and B cells respectively (**Supplementary Fig. 4a**), were both characterized by expression of upregulated ‘emergent’ genes such as *Atf5* and *Mthfd2* from the ISR pathway (30) (**Supplementary Fig. 4b, c** and **Supplementary Table 2**). These clusters also exhibited specific expression of other ISR pathway genes and an ISR gene module (30) identified in literature (**Fig. 3f**).

Cluster 2 cells, by contrast, were characterized by downregulation of ‘emergent’ genes such as *Pax2*, *Arvcf*, and *Shroom3* (**Supplementary Fig. 4d** and **Supplementary Table 2**). They also exhibited reduced expression of other PT developmental transcription factors and cell adhesion molecules such as *Maf* (35)*, Hnf4a* (36), and *Cdh2* (37) (**Fig. 3g**).

Taken together, these results indicate *Vhl* and *Pbrm1* inactivation regulate most genes in an additive manner. However, over time, their consequences converge to cause some *Vhl/Pbrm1*-null cells to adopt the stress response and lose expression of genes that confer epithelial identity and integrity.

### Maintenance of a *Vhl*-dependent proliferative drive following *Pbrm1* inactivation

Our transcriptional analysis did not highlight any *Pbrm1*-dependent changes in the expression of genes associated with the cell cycle. This was surprising considering the association between *PBRM1* inactivation and accelerated oncogenesis in ccRCC. We therefore studied the effects of *Pbrm1* inactivation on the survival and proliferation of RTE cells in further detail.

We first assessed the time-dependent changes in the number of cells bearing *Vhl* and/or *Pbrm1* inactivation by comparing the proportion of cells in different parts of the RTE that were positive for tdTomato across genotypes and timepoints. In the renal papilla, we had previously observed a HIF1A-dependent elimination of *Vhl*-null cells over time (18,20). Renal papillae from VPKO mice exhibited a decrease over time in the proportion of cells stained positive for tdTomato that was equivalent to the decrease seen in papillae from VKO mice over time. This suggested that *Pbrm1* inactivation did not rescue the elimination of *Vhl*-null cells in this non-cancer-associated tissue (**Fig. 4a** and **Supplementary Fig. 5a**).

**Fig. 4:**
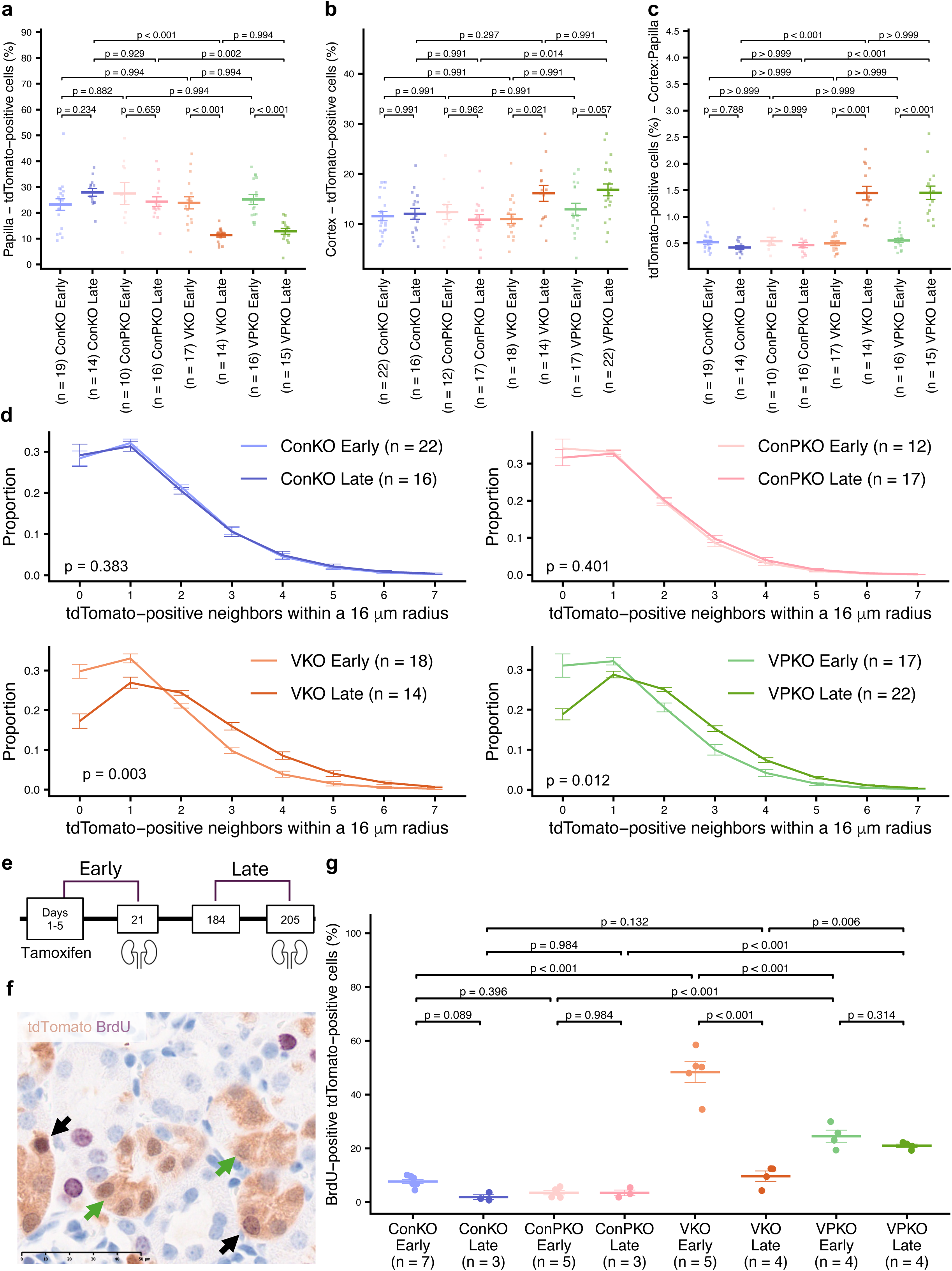
Maintenance of a *Vhl*-dependent proliferative drive following *Pbrm1* inactivation. **(a-c)** Automated quantification (see ‘**Methods**’) of the proportion of cells that are tdTomato-positive in the renal papilla **(a)** and cortex **(b)** of ConKO, ConPKO, VKO, and VPKO mice given 5x 2 mg tamoxifen and harvested early (1-3 weeks) and late (4-12 months) following recombination. The ratio between the proportion of cells that are tdTomato-positive in the cortex and the papilla **(c)** in the same mice. Error bars indicate Mean ± SE. Pairwise comparisons by two-tailed, two-way ANOVA test with multiple testing correction using the Benjamini-Hochberg method. The number of biological replicates (mice) analyzed for each genotype and timepoint is indicated. **(d)** Frequency distribution of tdTomato-positive neighbors of tdTomato-positive cells within a 16 μm radius (see ‘**Methods**’) in the cortex of ConKO, VKO, ConPKO and VPKO mice harvested early (1-3 weeks) or late (4-12 months) post recombination. Error bars denote Mean ± SE. Frequency distributions were compared using Wilks’ lambda statistic after multivariate analysis of variance (MANOVA) with Benjamini-Hochberg correction for multiple testing. **(e)** Outline of experimental protocols to measure BrdU incorporation in cells early or late after recombination. Harvest timepoints are depicted by the position of schematics of the kidneys. BrdU was administered continuously during a 21-day window immediately prior to harvest of cells performed either at an early (21 days) or late (205 days) timepoint. **(f)** Representative dual IHC for tdTomato (brown) and BrdU (purple) counterstained with hematoxylin. Examples of tdTomato-positive, BrdU-negative cells are depicted by green arrows; examples of tdTomato-positive BrdU-positive cells are depicted by black arrows. Scale bar, 50 μm. 40x magnification. **(g)** Manual quantification of the proportion of tdTomato-positive cells that are also BrdU-positive in the renal cortex of ConKO, ConPKO, VKO, and VPKO mice harvested early and late following recombination. Error bars indicated Mean ± SE. Pairwise comparisons were made by two-tailed, two-way ANOVA test with multiple testing correction using the Benjamini-Hochberg method. The number of biological replicates (mice) analyzed for each genotype and timepoint is indicated.

In the renal cortex, we had previously observed a HIF1A- and HIF2A-dependent accrual of *Vhl*-null cells over time (18,20). *Vhl/Pbrm1*-null cortical cells exhibited an increase in proportion over time that was equivalent to that of *Vhl*-null cells (**Fig. 4b** and **Supplementary Fig. 5b**). To account for variation in induced recombination between mice, we calculated tdTomato-positivity in the cortex as a ratio to tdTomato-positivity in the papilla of the same mouse. Expectedly, when treated this way, the time-dependent rise in the ratio was higher but also more consistent across mice, when compared to the analysis of the renal cortex on its own (**Fig. 4c**). This analysis confirmed that the time-dependent increase observed in VPKO mice was equivalent to that seen for VKO mice (**Fig. 4c**), indicative of tdTomato-positive cell proliferation and expansion along the renal tubule. To test this further, we reasoned that such cell proliferation should be reflected in a greater number of tdTomato-positive immediate neighbors at 4-12 months (late) versus 1-3 weeks (early) after recombination. As PT cells typically have at least one neighbor within an internuclear distance of 16 μm (20), we quantified tdTomato-positive neighbors of tdTomato-positive PT cells within this distance and compared over time across genotypes. This analysis also showed a time-dependent increase that was equivalent between the *Vhl*-null and *Vhl/Pbrm1*-null genotypes (**Fig. 4d**). These data indicated that additional *Pbrm1* inactivation did not alter the overall survival and accrual of *Vhl*-null PT cells between 1-3 weeks and 4-12 months following recombination in the ccRCC-prone renal cortex.

To study whether this equivalent accrual was derived from similar dynamics of cell proliferation, we administered the thymidine analog 5-bromo-2-deoxyuridine (BrdU) (38) to ConKO, VKO, ConPKO, and VPKO mice for a continuous period of 21 days immediately before harvest, either at 1-3 weeks (early) or 4-12 months (late) following recombination (**Fig. 4e**), and quantified the proportion of cortical tdTomato-positive PT cells that were positive for BrdU by dual immunohistochemistry (**Fig. 4f**). Consistent with prior work, an early proliferative drive was evident in *Vhl*-null cells (VKO Early versus ConKO Early) (18,20). This initial proliferative drive was present in *Vhl/Pbrm1*-null cells (VPKO Early versus ConPKO Early) but surprisingly was attenuated compared to *Vhl*-null cells (VPKO Early versus VKO Early) (**Fig. 4g**). Notably, while this drive was abrogated in *Vhl*-null cells over time (VKO Late versus VKO Early), it was maintained in *Vhl/Pbrm1*-null cells (VPKO Late versus VPKO Early) (**Fig. 4g**). *Pbrm1* inactivation alone did not result in either an initial (ConPKO Early versus ConKO Early) or time-dependent (ConPKO Late versus ConPKO Early) proliferation drive (**Fig. 4g**).

To test this further, we measured the proportion of cortical tdTomato-positive cells that were positive for the cell cycle marker Ki67 (39) by dual immunofluorescence microscopy (**Supplementary Fig. 5c**). This confirmed a strong but transient increase in cell cycling in *Vhl*-null cells and a modest but sustained increase in *Vhl/Pbrm1*-null cells **(Supplementary Fig. 5d**).

Taken together, these data indicated that *Pbrm1* inactivation did not alter the cell-type specific effects of *Vhl* inactivation on cell survival nor did it drive proliferation in cortical PT cells. However, it enabled cells to maintain a *Vhl*-inactivation-driven proliferative drive that is otherwise restrained in *Vhl*-null cells over time.

### Morphological alterations in *Vhl/Pbrm1*-null cells

Given our transcriptomic analyses highlighting early changes in genes involved in epithelial adhesion and morphogenesis, including downregulation of several epithelial polarity genes, as well as emergence of cell states characterized by a loss of genes contributing to polarity and tubular epithelial identity, we examined renal tubular morphology following *Vhl* and/or *Pbrm1* inactivation in detail. We visualized cells and studied their location within the tubular architecture by coupling tdTomato immunohistochemistry to Periodic Acid Schiff (PAS) staining. We first examined changes at 4-12 months (‘late’) following recombination, when our analyses had captured the emergence of cell states associated with a loss of epithelial identity and integrity.

PAS staining colors the basement membrane and the apical brush border of PT cells that surrounds the tubular lumen (40). In kidneys from VPKO mice harvested 4-12 months following recombination, we commonly observed (50-100 per section in n = 5/5 mice) tdTomato-positive cells that had escaped the tubular structure, either entering the lumen (‘luminal’ cells) or, less frequently (3-10 cells per section in n = 5/5 mice), protruding through the basement membrane into the interstitium (‘protruding’ cells) (**Fig. 5a**). Luminal tdTomato-positive cells detected by immunofluorescence microscopy had lost the expression of N-cadherin (CDH2), the canonical cell adhesion molecule in PT cells (**Fig. 5b**) (37). Protruding tdTomato-positive cells showed altered F-actin distribution, which is normally localized around the basolateral cortex and brush border of PT cells (**Fig. 5c**).

**Fig. 5:**
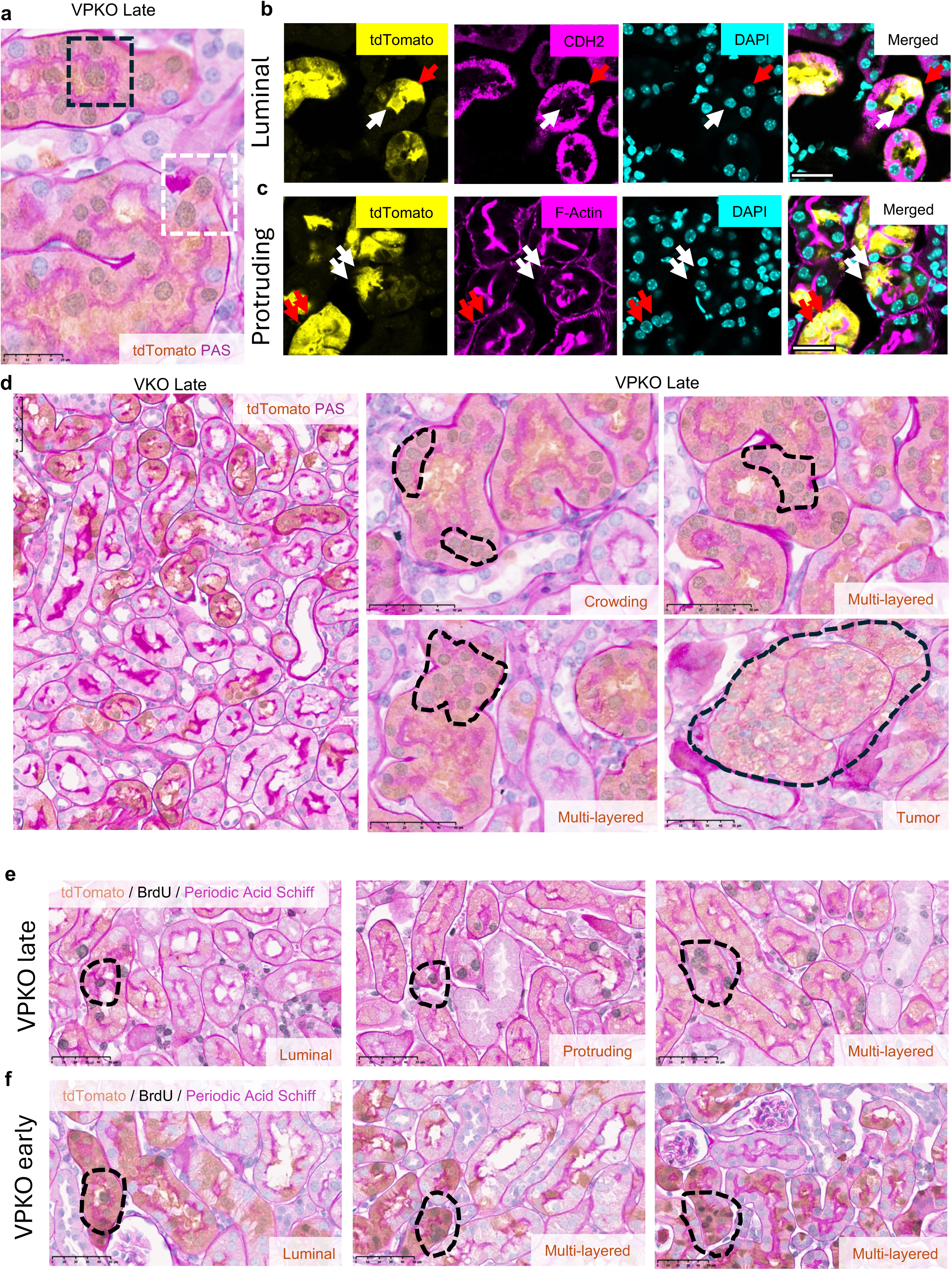
Morphological alterations in *Vhl/Pbrm1*-null cells. **(a)** Representative image of tdTomato IHC (brown) coupled with Periodic Acid–Schiff (PAS) staining (pink) of the renal cortex in VPKO mice (n = 5) harvested late (4-12 months) following recombination. PAS staining outlines the basement membrane and brush border of proximal tubules. Examples of tdTomato-positive cells that are located inside the tubular lumen (black dashed rectangle) or are protruding through the basement membrane into the renal interstitium (white dashed rectangle) are depicted. Scale bars, 25 μm. 40x magnification. **(b)** Representative images of immunofluorescence (IF) staining of tdTomato (yellow), N-cadherin (CDH2; magenta), and DAPI (cyan) in VPKO mice (n = 5) harvested late after recombination, showing an example of a ‘luminal’ tdTomato-positive cell that has lost CDH2 expression (white arrows), compared to a tubular tdTomato-positive cell that has retained CDH2 staining (red arrows). Scale bars, 25 μm. 20x magnification. **(c)** Representative images of IF staining of tdTomato (yellow), F-actin (magenta), and DAPI (cyan) in VPKO mice (n = 5) harvested late after recombination, showing examples of ‘protruding’ tdTomato-positive cells that have lost cortical F-actin localization (white arrows), compared to tubular tdTomato-positive cells that retain F-actin staining (red arrows). Scale bars, 25 μm. 20x magnification. **(d)** Representative tdTomato/PAS co-staining of renal cortex in VKO (n = 5) or VPKO mice (n = 5) harvested late (4-12 months) after recombination. Normal morphology is observed in the VKO mouse (left panel). In contrast, three types of morphological alterations are observed in kidneys from VPKO mice, including cell crowding within a monolayer (middle upper panel), multilayered epithelium (middle lower and right upper panels), and formation of tumors (right lower panel). Scale bars, 50 μm. 40x magnification. **(e, f)** Representative images of dual tdTomato (brown) and BrdU (grey) IHC coupled to PAS staining (pink) performed in renal sections from VPKO mice harvested late (n = 5) (**e**) or early (n = 5) (**f**) following recombination and counterstained with hematoxylin. Examples of luminal, protruding, and multilayered tdTomato-positive cells that have incorporated BrdU are denoted by dashed black circles. Scale bars, 50 μm. 40x magnification.

PT cells are normally organized in a single layer, spaced evenly around the tubular circumference. This morphology was retained in kidneys from VKO (n = 5/5) mice harvested 4-12 months (late) following recombination (**Fig. 5d**). However, in VPKO mice, we commonly observed (20-40 instances per section in n = 5/5 mice) regions where nuclei of neighboring tdTomato-positive cells had crowded together within the monolayer (**Fig. 5d**). Moreover, tdTomato-positive cells also formed multi-layered clusters within the tubule (**Fig. 5d**). These clusters ranged from small groupings of 5-10 cells (10-15 clusters per section in n = 5/5 mice) and extended to small tumors of up to 50 cells in a single section (2-5 tumors detected across 30 sections stained in n = 2/5 mice), uniformly stained for tdTomato (**Fig. 5d**). In such clusters and tumors, a luminal brush border was not visible, but some cells exhibited circumferential, as opposed to strictly apical and basal, PAS staining that was suggestive of loss of polarity (**Fig. 5d**).

We wondered if the multi-layered clusters and small tumors in kidneys of VPKO mice reflected unrestrained proliferation of *Vhl/Pbrm1*-null cells. To test this, we administered BrdU continuously for 21 days before harvest and studied its incorporation using dual tdTomato/BrdU immunohistochemistry that was coupled to PAS staining. In kidneys from ConKO and VKO mice harvested 4-12 months (late) following recombination, tdTomato-positive BrdU-positive cells remained within the confines of the proximal tubule and were regularly spaced in a monolayer. In contrast, in VPKO mice at a similar late timepoint, tdTomato-positive BrdU-positive cells often broke the tubular boundary, were frequently crowded along the tubular monolayer, and formed multi-layered cell clusters (**Fig. 5e**).

Finally, to resolve whether these abnormalities were a direct result of proliferation or instead due to a ‘release’ from epithelial constraints caused by *Pbrm1* inactivation, we analyzed BrdU-positive, tdTomato-positive cells in ConPKO mice at 4-12 months (late) following recombination, and in VKO and VPKO mice at 1-3 weeks (early) following recombination. ConPKO mice had far fewer BrdU-positive, tdTomato-positive cells than VPKO mice at a late timepoint, but these were often luminal or crowded along the tubular circumference, suggesting such abnormalities arise even without a sustained proliferative drive (**Supplementary Fig. 6a**). Similarly, luminal, crowded or multi-layered BrdU-positive, tdTomato-positive cells were far more frequent in kidneys of VPKO (**Fig. 5f**) compared to VKO mice (**Supplementary Fig. 6b**) at an early timepoint, despite the latter containing more proliferating cells.

Taken together, these analyses suggested that through an early and *Vhl*-independent action, *Pbrm1* inactivation disrupts epithelial organization of *Vhl/Pbrm1*-null PT cells. This allows cells to breach the tubular structure, form multi-layered epithelia, retain proliferative capacity, and eventually form renal tumors.

## Discussion

In this work we sought to understand the oncogenic interplay between inactivation of *VHL* and *PBRM1*, the most common genetic interaction in the pathogenesis of ccRCC. In humans, combined biallelic inactivation of *VHL* and *PBRM1* is associated with accelerated progression towards ccRCC (6). This can be modelled in the mouse, where combined inactivation of *Vh*l and *Pbrm1* but not inactivation of *Vhl* alone, results in the development of tumors in the renal cortex that mimic many aspects of the human disease (8–10). To better understand this process, we deployed an oncogenic cell tagging system that tightly couples the inactivation of *Vhl* to the expression of tdTomato, enabling study of the earliest events in the progression to renal tumor formation. This model enabled us to retrieve *Vhl*/*Pbrm1*-inactivated marked cells from the native kidney, examine changes in gene expression at cell resolution, follow them in time and relate them to phenotypic effects including the proliferation and accrual of marked cells and the development of morphological abnormalities.

Although it has been proposed that the VHL-PBRM1 interaction serves to modulate the HIF transcriptional pathway that is activated when VHL is defective (10,14,16), we found no evidence for this here. Specifically, we found no significant associations between genes regulated by *Pbrm1* and genes regulated by hypoxia or HIF, as defined in different published reports, including lists of HIF target genes defined previously in this model (20,25). scRNA-seq analyses also allowed for testing for such associations in specific classes of the proximal renal tubule and at different times after gene inactivation. Again, we observed no association, either positive or negative between HIF target genes and those whose expression was altered by *Pbrm1* inactivation. Rather we found that the effects of *Pbrm1* inactivation were largely distinct, both from effects of activation of either HIF1A or HIF2A, and from the overall effects of *Vhl* inactivation itself. This was true when gene expression patterns were compared between *Vhl*- and combined *Vhl*/*Pbrm1*-inactivation both early and late after recombination.

Our findings do not exclude effects of PBRM1 on HIF under other contexts and therefore are not necessarily in conflict with the existing literature. However, the model we studied defines both the early and later consequences of the VHL/PBRM1 interaction on gene expression and develops tumors resembling ccRCC. Hence, we conclude that global modulation of the HIF transcriptional pathways is not necessary for PBRM1-associated oncogenesis.

In an attempt to define other interactions between VHL and PBRM1 at the level of gene expression, we also tested for an excess of observed changes in gene expression in cells retrieved late (4-12 months) after combined *Vhl/Pbrm1* inactivation compared to changes calculated from the sum of individual measures of *Vhl*-dependent and *Pbrm1*-dependent changes. This revealed a small group of ‘emergent’ genes, raising a question as to whether they drive, or are a consequence of, oncogenesis. The findings that several have been identified as ‘stress’ genes, that epithelial stress has been identified as a consequence of *Pbrm1* inactivation in other settings (15) and that they appeared to be expressed in specific groups of cells rather than generally across all cells bearing combined *Vhl/Pbrm1*-inactivation, suggests that they are secondary to the abnormal epithelial state as opposed to being drivers of oncogenesis.

Among gene expression changes observed following *Pbrm1* inactivation were those with functions in lipid metabolism. Consistent with this we found that cells lacking *Pbrm1* manifest cytoplasmic lipid droplets, a common but poorly understood finding in cancer including ccRCC (41). However, of particular interest was the over-representation of genes with functions in epithelial morphogenesis, integrity and cell adhesion. These findings were similar whether one (ConPKO) or both (VPKO) *Vhl* alleles were inactivated, consistent with independence from HIF. In keeping with this, similar enrichment of *Pbrm1-*regulated genes within GO terms for such processes has been observed in other, non-VHL associated types of cancer such as breast cancer (15), where HIF is not activated genetically.

A key aim of the current work was to combine these genotypic analyses with concurrent morphological analyses enabled by the oncogenic cell tagging strategy and to focus on the nature and timing of the earliest changes that could illuminate the VHL-PBRM1 interaction. Although *Vhl* is a classical tumor suppressor, with biallelic inactivation being observed as the common truncal genetic event in the pathogenesis of ccRCC, in most contexts, inactivation of *Vhl* results paradoxically in an anti-proliferative phenotype (42–44). One exception to this is the renal tubule where in the intact mouse kidney, proliferation of proximal renal tubular cells is observed early after *Vhl* inactivation, which is dependent on the integrity of both HIF1A and HIF2A (18,20). Remarkably, examination of this phenotype using the oncogenic cell tagging strategy revealed that in *Vhl*-inactivated kidneys, this proliferation, although initially marked, is transient (18,20). Interestingly, it led to an accumulation of tdTomato positive cells in renal tubules, but the cells appeared to be strictly restrained within the boundaries of the normal renal tubular epithelial monolayer.

Using the oncogenic cell tagging system we examined changes in cell proliferation, cell morphology and tubular integrity in further detail and directly compared observations following *Vhl* inactivation alone with those of combined *Vhl* and *Pbrm1* inactivation. Surprisingly, when compared to the effects of inactivation of *Vhl* alone, this revealed that indices of proliferation were not increased and were even reduced early after the combined inactivation of *Vhl* and *Pbrm1*. However, in contrast to inactivation of *Vhl* alone, proliferation was sustained. In association with this, we observed a series of abnormalities compromising the integrity of the renal tubular epithelium, including loss of N-cadherin, dysregulation of F-actin, and epithelial cell crowding. Strikingly, cells were observed moving out of the renal tubular epithelial monolayer into the tubular lumen, or into the renal interstitium or forming a multi-layered epithelium. Epithelial abnormalities were present early after combined *Vhl* and *Pbrm1* inactivation but were more extensive at later time points where they were accompanied by multi-focal tumor formation. Based on the combined cell biological and genetic data, we propose that the necessary and sufficient oncogenic interaction of VHL with PBRM1 is integrated not at the level of effects on gene expression but through distinct cell biological effects on the renal epithelium that, when combined, result in dysregulated growth.

Chromatin immunoprecipitation has defined approximately 50,000 PBRM1 genome binding sites, selected through combinatorial binding of bromodomains to acetylated lysine residues on histone tails (45,46). Other potentially relevant physical connections have been defined between methylated lysine on tubulin and the BAH domains of PBRM1 (13). Although our work suggests that changes in the integrity of the renal tubules consequent on PBRM1 inactivation are a key component of PBRM1-associated oncogenesis, further work will be needed to define the precise mechanisms linking PBRM1 integrity with epithelial function. In that context, the complete gene expression datasets associated with this work are provided as a resource.

Nevertheless, our work provides a new conceptual framework for understanding the oncogenic interaction of VHL and PBRM1 inactivation in kidney cells, which we hypothesize could be of general importance in cancer. Defining the interface between anatomical integrity and control of cell proliferation is arguably the central problem in understanding physiological growth and its dysregulation in cancer. In most forms of cancer, it is assumed that increased cell proliferation drives dysregulation of structure, but it is difficult to distinguish cause and effect. Does dysregulated proliferation drive structural dysregulation or does dysregulation of structure release restraints on cell proliferation? The ability to observe cell behavior before and after the development of morphological abnormalities and to relate this to patterns of gene expression provided a new opportunity to dissect these questions. Our data suggest that *Vhl* inactivation results in HIF-dependent proliferative drive within proximal tubular cells that is ordinarily restrained by systems that maintain the anatomical integrity of the renal tubular epithelium. We propose that, at least in this context, rather than directly increasing proliferative drive, *Pbrm1* inactivation removes or impairs that restraint, permitting continued proliferation of cells outside the confines of the tubular epithelial monolayer. Thus, combined inactivation of *Pbrm1* and *Vhl* results in progressive epithelial dysplasia and ccRCC-like tumor formation. Given the extensive association of mutations in PBRM1 with other tumors (47), this model may be relevant to a broader range of epithelial cancers.

## Methods

### Mice

All experimental procedures were conducted following approval by the Medical Science Ethical Review Committee of the University of Oxford and authorized under UK Home Office regulations of Animals (Scientific Procedures) Act 1986.

Mice were housed in individually ventilated cages on a 13 hr light/ 11 hr dark cycle with food and water provided *ad libitum*. B6.*Vhl^tm1.1b(tdTomato)Pjr^* (*Vhl^pjr.fl^*) mice, which allow for conditional excision of *Vhl* exons 2 and 3 along with simultaneous tdTomato insertion, were commissioned from Ozgene, Australia and generated using goGermline technology (48). *Vhl^tm1jae^* mice (23) (*Vhl^jae.fl^*) (RRID: IMSR_JAX:012933), were crossed with Tg(Pgk1-cre)1Lni (49) (*Pgk1-cre*) (RRID: IMSR_JAX:020811) to generate a constitutively inactivated *Vhl^tm1jae^* allele (*Vhl^jae.KO^*), in which the *Vhl* promoter and exon 1 are excised. Tg(Pax8-cre/ERT2)CAmat(24) (*Pax8-CreERT2*) (RRID: IMSR_HAR:9175) mice were obtained *via* the European Mouse Mutant Archive (EMMA). Mice carrying conditional alleles for *Pbrm1* (*Pbrm1^tm1c(EUCOMM)Wtsi^*; RRID:IMSR_JAX:031875; termed *Pbrm1^fl^*) (8), which allow for conditional excision of *Pbrm1* exon 2, were obtained from JAX.

Mice of both sexes, weighing at least 20 g, were administered tamoxifen by oral gavage daily for five consecutive days at a dose of 100 μl of 20 mg/ml tamoxifen (Sigma T5648) in 10% (v/v) ethanol in corn oil (Sigma C8267).

BrdU was administered to mice through drinking water. 1 mg/ml BrdU (Sigma B5002) was prepared in 1% (w/v) sucrose solution in double-distilled water and provided *ad libitum* for a period of 21 days before harvest. BrdU water was protected from light and replaced every week.

The genotypes of mice bred, the expected *Vhl* and *Pbrm1* status of cells before and after Cre-mediated recombination, and abbreviations used in this paper are detailed in **Table 1**. Mice were harvested either 1-3 weeks (termed ‘early’ timepoint), 4-12 months (termed ‘late’ timepoint), or 12-17 months (termed ‘aged’ timepoint) after recombination was induced.

### Genotyping and PCR

Genotypes of all experimental mice were confirmed at time of harvest. Primer sequences and expected product sizes were as follows: *Vhl^jae.KO^*: 5’-CTGGTACCCACGAAACTGTC-3’, 5’-CTAGGCACCGAGCTTAGAGGTTTGCG-3’ and 5’-CTGACTTCCACTGATGCTTGTCACAG-3’ (260 bp for *Vhl^wt^,* 260 and 240 bp for *Vhl^jae.KO^* multiplex products); *Vhl^pjr.fl^*: 5’-GGTGCTAATTGAAGGAAGCTACTG-3’ and 5’-CTCCTCCGAGGACAACAACATG-3’ (1067 bp); *Pax8-CreERT2*: 5’-CGGTCGATGCAACGAGTGATGAGG-3’, 5’-CCAGAGACGGAAATCCATCGCTCG-3’, 5’-CTCATACCAGTTCGCACAGGCGGC-3’ and 5’- CCGCTAGCACTCACGTTGGTAGGC-3’ (300 bp for wild-type, 300 bp and 600 bp multiplex products for transgenic); *Pbrm1^fl^*: 5’-ATCTCTCCAGACCCCCAAAC-3’ and 5’-AAGACCCTGCCTCAAAACAC-3’ (243 bp for *Pbrm1^wt^*, 400 bp for *Pbrm1^fl^*).

### Immuno-labelled and histological staining

#### Tissue preparation

Mice were euthanized with isoflurane anesthesia, flushed via the aorta with 1x PBS pH 7.4 (Gibco 70011) to clear blood from tissues, and perfused with 4% (w/v) paraformaldehyde (Sigma P1213) in PBS at room temperature (RT). Kidneys were bisected and fixed in 10% (v/v) neutral buffered formalin (NBF; Sigma HT501128) with rocking for 24 hr at RT. Kidneys were then dehydrated through a graded ethanol series (70% to 100%) and xylene before paraffin embedding. Embedded tissues were cut to 4 μm sections on a Thermo Microm HM 355S Microtome using MB35 Premier Blades. Sections were floated on warm distilled water and mounted on poly-lysine coated slides (Fisher 10149870). Slides were dried for at least 3 hr at 37°C before immunohistochemistry (IHC).

For frozen sections, mice were euthanized with terminal isoflurane anesthesia, flushed via the aorta with ice-cold 1x PBS pH 7.4 (Gibco 70011) to clear blood from tissues, kidneys were bisected and fixed with ice-cold 4% (w/v) paraformaldehyde (Sigma P1213) in PBS pH 7.4 for 24 hr. To protect tissues from freezing artefacts, fixed kidneys were immersed in ice-cold 30% (w/v) sucrose in PBS pH 7.4 overnight, then embedded in optimum cutting temperature (OCT) compound (Tissue Tek). Kidneys were cut to 8-10 μm sections at -18°C on a Bright OTF700 Cryostat and mounted on poly-lysine coated slides.

#### Hematoxylin and Eosin (HE) Staining

Formalin-fixed paraffin embedded (FFPE) sections were deparaffinized with xylene and ethanol, rehydrated with double-distilled water, and immersed in modified Harris Hematoxylin (Fisher 72711) for 3 min. Staining was differentiated by dipping slides in 0.25% (v/v) HCl in 70% ethanol for 10 s, washing twice in tap water. Hematoxylin staining was blued by immersion in 0.06% (v/v) NH_4_OH in water for 30 s. Slides were then washed in 90% and 100% ethanol for 30 s and transferred to Shandon Alcoholic Eosin Y solution (Fisher 6766008) for 3 min. Slides were washed in 100% ethanol, dehydrated with xylene, and mounted with DPX mountant (Merck 06522).

#### HIF1A and HIF2A IHC

For the detection of HIF1A and HIF2A, a new signal amplification system was developed in the laboratory. FFPE kidney sections were deparaffinized, rehydrated and subjected to heat-induced epitope retrieval (HIER) with the Dako Target Retrieval Solution pH 6.0 (S2369, Agilent) for 30 min in a pressure cooker (Tefal; set to 170 kPa). Endogenous peroxidase activity was blocked by incubation with Dako Peroxidase Blocking solution (Agilent S2023) for 10 min at RT. Non-specific protein binding was blocked with 5% (w/v) bovine serum albumin (BSA; Sigma 5482) in 1x TBS-T (50 mM Tris, 31.6 mM NaCl, 0.1% (v/v) Tween-20; pH 8.4) and 10% (v/v) goat serum for 40 min at RT. Endogenous avidin and biotin activity was blocked with the Avidin and Biotin blocking kit (Abcam ab64212) for 5 min each and slides were incubated with anti-HIF-1α primary antibody (Cayman 1000642; 1:1000) and anti-HIF-2α primary antibody (PM8 (50); 1:3000) overnight at 4°C. Indirect labelling was performed by serial incubations with secondary HRP-conjugated anti-rabbit antibody (Dako K4003, Agilent; 30 min RT), tertiary biotin-conjugated anti-HRP antibody (Rockland 200-4638-0100; 1:250; 2 hr RT) and quaternary HRP-conjugated Streptavidin (BD Pharmingen 551321; 1 hr RT). Signal was detected with the Dako Envision system (Agilent K4003), visualized using diaminobenzidine (DAB) and counterstained with modified Harris Hematoxylin (Fisher 72711) differentiated for 10 s in 0.25% (v/v) HCl in 70% ethanol. Hematoxylin staining was blued by immersion in 0.06% (v/v) NH_4_OH in water for 30 s. Slides were dehydrated and mounted with DPX mountant (Merck 06522).

#### tdTomato and PBRM1 IHC

Sections were deparaffinized with xylene and ethanol, rehydrated with double-distilled water, and subjected to HIER with TE buffer (10 mM Tris, 1 mM EDTA, pH 9.0) in a steamer (Braun S900087) for 20 min for tdTomato IHC or with the Dako Target Retrieval Solution pH 6.0 (S2369, Agilent) for 10 min in a pressure cooker (Tefal; set to 170 kPa) for PBRM1 IHC. Endogenous peroxidase activity was blocked by incubation with Dako Peroxidase Blocking solution (Agilent S2023) for 10 min at RT. Non-specific protein binding was blocked by incubation with 5% (w/v) bovine serum albumin (BSA; Sigma 5482) in 1x TBS-T (50 mM Tris, 31.6 mM NaCl, 0.1% (v/v) Tween-20; pH 8.4) for 40 min at RT. Antigens were detected by overnight incubation at 4°C with primary antibodies diluted 1:1000 in Dako Antibody Diluent Solution (Agilent S3022): tdTomato – anti-RFP (Rockland 600-401-379, RRID:AB_2209751); PBRM1 – anti-PBRM1 (Proteintech 12563-1-AP, RRID:AB_2877865).

Slides were washed in 1x TBS-T. Signal was detected with the Dako Envision system (Agilent K4003), visualized using diaminobenzidine (DAB), and counterstained with modified Harris Hematoxylin (Fisher 72711) differentiated for 10 s in 0.25% (v/v) HCl in 70% ethanol. Hematoxylin staining was blued by immersion in 0.06% (v/v) NH_4_OH in water for 30 s. Slides were dehydrated and mounted with DPX mountant (Merck 06522).

#### tdTomato-BrdU Dual IHC

FFPE slides were deparaffinized and rehydrated as above and were subjected to HIER with TE buffer (10 mM Tris, 1 mM EDTA, pH 9.0) in a steamer (Braun S900087) for 20 min before proceeding with tdTomato IHC as described above. After DAB exposure (brown in color), slides were rinsed in distilled water, and any remaining peroxidase activity was blocked by immersing slides in 3% (w/v) H_2_O_2_ in PBS pH 7.4 for 10 min at RT. BrdU was detected using the BrdU Detection Kit II (BD 551221) following manufacturer’s protocols. Slides were washed in PBS and subjected to HIER with the antigen retrieval reagents provided in the BrdU Detection Kit II (BD 551221) for 10 min in a pressure cooker (Tefal; set to 170 kPa). After slides had cooled down, they were counterstained with hematoxylin and blued as described above. Counterstained slides were then rinsed in distilled water and subjected to endogenous avidin and biotin activity blocking, and non-specific protein binding blocking with the Avidin and Biotin Blocking kit (Abcam ab64212) and DAKO protein Block (Agilent X0909) respectively for 10 min each at RT. Slides were incubated with anti-BrdU antibody for 2 hr at RT. Signal was generated by incubation with streptavidin-HRP (BD Biosciences 550946) at RT for 30 min and visualized using the Vector VIP (purple in color) substrate (Vector SK4605), which contrasted with the DAB (brown) signal produced previously on the same sections for tdTomato. Slides were dehydrated by quick dips in 100% ethanol and xylene and were mounted with DPX mountant (Merck 06522).

#### tdTomato and tdTomato/BrdU IHC coupled with PAS Staining

PAS staining colors the basement membrane and the apical brush border of PT cells that surrounds the tubular lumen (40), allowing for careful analysis of renal tubular anatomy and structure. For tdTomato IHC coupled to PAS staining, tdTomato IHC was performed first as described above. tdTomato signal was visualized after a brief 30 s exposure to DAB (brown) and slides were rinsed in distilled water. Slides were then immersed in 0.5% Periodic Acid Solution (Sigma 1004821000) for 15 min at RT and washed four times in distilled water. Next, slides were immersed in Schiff’s Reagent (Sigma 3952016) for up to 2 min with heating in a microwave (Sharp; set to 900 W), until the solution was colored dark pink. Subsequently, slides were rinsed in running hot tap water until the pink solution had completely cleared. Slides were then counterstained with hematoxylin and blued as described above, before dehydration and mounting in DPX.

For tdTomato-BrdU dual IHC coupled to PAS staining, tdTomato IHC was performed as described above for tdTomato-PAS staining. BrdU detection followed as described above, but without the interceding hematoxylin counterstaining step. Here, BrdU signal was visualized with the ImmPACT SG (grey) substrate (Vector SK4705), which contrasted with the DAB (brown) stain used to detect tdTomato on the same sections. After tdTomato and BrdU detection, PAS staining and hematoxylin counterstaining were performed as described above.

#### Immunofluorescence (IF)

Frozen sections were brought to RT and washed with PBS for 10 min at RT. They were permeabilized with 0.2% (v/v) Triton-X (Sigma X100). Endogenous avidin and biotin activity was blocked with the Avidin and Biotin Blocking kit (Abcam ab64212) for 5 min each at RT. Non-specific protein binding was blocked with 5% (w/v) BSA 10% (v/v) donkey serum in TBS-T for 1 hr at RT. Sections were incubated overnight at 4°C with the following primary antibodies, all diluted 1:100 in the blocking solution: Ki67 (Invitrogen 13-5698-82 (biotinylated), RRID:AB_2572794), N-cadherin (Novus NBP1-48309B (biotinylated), RRID:AB_11039634), tdTomato (Rockland 600-401-379, RRID:AB_2209751). Slides were washed in TBS-T and signal detected with Alexa Fluor-labelled secondary antibodies or streptavidin (for tdTomato: Thermo A32795TR, RRID:AB_2866494; for Ki67 and N-Cadherin: Thermo S21374, RRID:AB_2336066) for 30 min at RT. F-Actin was detected directly by incubating sections for 30 min at RT with Alexa-Fluor 647 conjugated Phalloidin (Thermo A22287, RRID:AB_2620155) diluted 1:5000 in the blocking solution. Slides were washed and counterstained with 1 μg/ml 4’,6-diamidino-2-phenylindole (DAPI; Sigma D9542) for 5 min at RT and mounted with Vectashield Vibrance Antifade Mounting Medium (Vector H-1700).

#### Oil Red O staining

Frozen sections were brought to RT and washed with PBS for 10 min at RT. Staining was performed using the Oil Red O Stain kit (Abcam ab150678) following manufacturer’s instructions. Slides were placed in propylene glycol for 5 min and incubated in heated (60°C)

Oil Red O Solution for 15 min. Slides were washed in 85% fresh propylene glycol in distilled water for 1 min and rinsed in two changes of distilled water. Slides were counterstained with hematoxylin as described above and mounted in Vectashield Vibrance Antifade Mounting Medium (Vector H-1700).

#### Microscopy

IHC, PAS, and Oil Red O-stained sections were scanned with a Hamamatsu NanoZoomer S210 slide scanner at 40x magnification and analyzed with Hamamatsu NDP.view2 software. IF sections were scanned using the ‘tiling’ feature on a Zeiss LSM980 confocal microscope at 20x magnification. The laser intensity and pinhole settings were kept uniform for each combination of antibodies across all sections.

#### IHC and IF Quantification

The proportion of cells positive for tdTomato by IHC was quantified using the HALO Image Analysis Software (v3.5 and v3.6; Indica Lab, Albuquerque, USA). Kidney sections were annotated manually to define regions of the renal papilla or the renal cortex based on morphological appearance and anatomical location in the tissue (51). Briefly, the renal cortex was identified by the presence of glomeruli and proximal convoluted tubules (corresponding to PT S1 and S2 cell types), distal convoluted tubules, and cortical collecting ducts, each of which have characteristic tubular shape, nuclear shape, and nuclear distribution along the tubular circumference. The renal papilla, comprising cells of the collecting duct and located at the apex of the medullary pyramid, was characterized by its elongated tubules and sparse cellularity, giving it a relatively pale appearance compared to the cortex and outer medulla. Nuclei were detected using the HALO AI v3.6 Nuclei Seq algorithm based on hematoxylin counterstaining. Nuclei <15 µm^2^ and >200 µm^2^ were excluded from analysis. The Red-Green-Blue (RGB) IHC image was deconvoluted computationally into individual contributions from DAB or hematoxylin based on known colorimetric properties of the two stains. Cells were recorded as tdTomato-positive if the tdTomato signal passed a specified intensity threshold in detected nuclei and in a 2 µm perimeter drawn around each nucleus.

The proportion of tdTomato-positive cells that were also positive for Ki67 was quantified using the QuPath (v0.6.0) image analysis software (52). The kidney cortex was manually annotated on the scanned slides using the criteria listed above. Nuclei were detected on the basis of DAPI staining using QuPath’s ‘Cell Detection’ plugin. Nuclei <15 µm^2^ and >200 µm^2^ were excluded from analysis. As recommended by QuPath’s developers in the introductory vignette, true cellular signal was distinguished from tissue autofluorescence by ‘smoothing’, i.e. calculating the fluorescence intensity of each stain in a cell as a weighted average over a 25 μm radius neighbourhood of the cell. Cells were recorded as tdTomato-positive if the weighted tdTomato signal passed a fixed intensity threshold in detected nuclei and in a 2 µm perimeter drawn around each nucleus. Cells were scored as positive for Ki67 if the weighted Ki67 signal passed a fixed intensity threshold in the nucleus.

The proportion of cortical tdTomato-positive cells that were also positive for BrdU was quantified manually. Five 10x images, each covering 1.1 mm^2^ of cortical tissue, were captured from slide scans of tdTomato/BrdU dual IHC performed across mice of different genotypes and timepoints. The images were then ordered randomly and anonymized so that researchers were blind to mouse identity and genotype when counting.

Cells were identified as being ‘luminal’, ‘protruding’, ‘crowded’, or ‘multi-layered’ by assessing the position of their tdTomato-positive nuclei relative to the tubular lumen and the tubular basement membrane that were stained directly by PAS. Because of the convoluted nature of the proximal tubule, it is possible to mistake a cell that is part of the tubular structure but outside the plane of sectioning, for a cell that is mis-located within the tubular structure. Therefore, to ensure accurate identification of such cells, we first assessed the changing shape and orientation of proximal tubules by examining serial sections and then evaluated whether any tdTomato-labelled nuclei were aberrantly located. Through this exercise, we noted that whereas cells that appeared to be outside the tubular structure but were merely outside the plane of sectioning often stained uniformly and intensely with PAS, cells that were truly luminal or protruding were often surrounded by discontinuous or diffuse PAS staining and exhibited a shift perpendicular to the orientation of the tubule. ‘Crowded’ cells were identified as PT cells whose nuclei often contacted or overlapped one another in a single layer along the tubular circumference. When such cells formed multiple layers between the PAS-stained lumen and the basement membrane, they were identified as ‘multi-layered’.

### Tissue Dissociation

Kidneys were bisected, the renal capsule removed and then macerated for up to 7 min on a bed of ice. Macerated kidneys were then subjected to single-cell dissociation using the Multi-Tissue Dissociation Kit 2 (Miltenyi 130-110-203). Briefly, macerated tissues were suspended in 1.45 ml Buffer X, 30 µl of Enzyme D, 15 µl of Enzyme P, 15 µl of Buffer Y, and 6 µl of Enzyme A, all prepared according to the manufacturer’s instructions. The suspension was then incubated in a water bath at 37°C for 30 min in a shaking incubator set to 150 rpm. The procedure was stopped by the addition of 150 µl fetal bovine serum (FBS; Sigma F7524) and resuspended in 9 ml of RPMI-1640 medium (Merck R0883). The digest was filtered through a 40 µm cell strainer to remove undigested tissue and the filtrate was centrifuged (300g for 10 min at 4°C). Erythrocytes were eliminated by resuspending dissociated cells in 3 ml of 1x RBC Lysis Buffer (Miltenyi 130-094-183) prepared in deionized water and incubating for 2 min at RT. Cells were centrifuged (300g for 5 min at 4°C) and resuspended in ice-cold D-PBS before being counted on a Thermo Scientific Countess II machine for total yield and viability.

### Fluorescence Activated Cell Sorting

Dissociated cells were resuspended in 10% FBS, 2 mM EDTA, in D-PBS (Thermo 14190144) to a concentration of 5,000,000 live cells per ml. DAPI (Sigma D9542) was added to stain for viability. Cells were sorted using a BD Aria Fusion Cell Sorter. tdTomato was excited with a 561 nm laser and fluorescence detected with a 582/15 band pass filter. DAPI was excited with a 405 nm laser and fluorescence detected with a 450/40 band pass filter. The gating strategy is provided in **Supplementary Fig. 1b**. Live, single, tdTomato-positive cells were collected in FBS-coated polypropylene tubes and pelleted by centrifugation at 300g for 10 min. Cells were counted again for yield and viability and processed for DNA extraction, immunoblotting, or scRNA-seq.

### Immunoblotting

Protein extraction was performed by solubilizing cell pellets recovered from FACS and frozen over dry ice in Urea SDS buffer (7M Urea, 10 mM Tris, 1% SDS, 10% glycerol, pH 7.4), adding Laemmli Sample buffer to 1.5x (v/v) and a final concentration of 7,000 cells/μl, and then boiling at 95°C for 5 min. Immunoblotting was performed as described previously (53). Briefly, proteins were separated by SDS-polyacrylamide gel electrophoresis, transferred to polyvinylidene difluoride membrane (Immobilon-P, Millipore IPVH00010), and blocked in 4% (w/v) fat free milk in 1x PBS with 0.1% Tween 20. Primary antibodies: tdTomato (Rockland 600-401-379, RRID:AB_2209751) and PBRM1 (Proteintech 12563-1-AP, RRID:AB_2877865) were diluted in blocking buffer (1:1,000). HRP-conjugated secondary antibodies (Agilent P0448, RRID:AB_2617138) and chemiluminescence substrate (West Dura, Thermo Fisher Scientific 34076) were used to visualize proteins, using a ChemiDoc XRS+ imaging system (BioRad). After immunoblot analysis, membranes were stained with Coomassie brilliant blue to visualize separated proteins, and this was used as a reference of sample loading.

### Single-cell Library Preparation and Sequencing

Sorted cells were prepared into single-cell droplets using the Chromium Next GEM Single-cell kit. 10,000 live cells per sample were loaded on separate Chromium Next GEM Chip G (10X Genomics PN-1000127) channels. cDNA clean-up, amplification and adaptor ligation were performed with the Chromium Next GEM Single Cell 3ʹ Kit v3.1 (10X Genomics PN-1000268). cDNA yield was quantified using the High Sensitivity D1000 ScreenTape Assay (Agilent 5067-5584 and 5067-5585) to optimize the number of reaction cycles for library preparation. Single cell sequencing libraries were prepared using the Library Construction Kit (10X PN-1000190) and Dual Index Kit TT Set A (10X Genomics PN-1000215) sequencing indices.

Single-cell libraries were sequenced on an Illumina NextSeq 2000 sequencing system with P3 200 cycle reagents (Illumina 20040560). Libraries from up to six samples were pooled in an equimolar ratio, diluted to 650 pM in RSB buffer and mixed with 1% PhiX Control v3 DNA (Illumina FC-110-3001) for sequencing. Preliminary sequencing results (bcl files) and FASTQ files were generated with the DRAGEN BCL Convert workflow, using the workflows optimized for 10X Chromium 3’ v3.1 Dual Index gene libraries, with 28 and 152 Read 1 and 2 cycles respectively.

### Single-Cell Sequencing Data Processing and Quality Control

The mm10 reference genome was downloaded from https://cf.10xgenomics.com/supp/cell-exp/refdata-gex-mm10-2020-A.tar.gz. The transcript sequence and annotation for *Vhl^pjr.KO^,* that included the tdTomato transgene, were added manually to the FASTA reference genome file and GTF file respectively using the CellRanger (v6.1.1) mkref function. FASTQ files generated for each sample were aligned to the custom reference genome by CellRanger using the default parameters. After aligning, for each read pair, cell barcodes and unique molecular identifiers (UMIs) were obtained from Read 1 and read counts per feature were obtained from Read 2. Only those UMIs that could be linked to a valid cell barcode and a gene exon region were included to create the cell by gene count matrix. Reads from *Vhl* exon 1 were excluded from analysis to prevent ambiguous alignment to *Vhl^wt^* or *Vhl^pjr.KO^* alleles. Cells were subjected to the following filters: detected genes per cell > 200, fraction of mitochondrially-encoded reads per cell < 0.5 and detected genes < 3x median for each sample. The threshold for mitochondrially encoded reads was set to this value in line with published kidney scRNA-seq studies to account for the high mitochondrial content in the RTE cells (54). Data from cells from different samples and sequencing runs were then aggregated, which sometimes required manual renaming of duplicate cell ‘names’. Downstream analysis was conducted using the R package Seurat v5.1.0 (55).

### Dimension Reduction and Clustering

Gene expression in PT cells is known to exhibit sex-specific differences (56,57). To account for these during clustering and dimension reduction analysis, we randomly downsampled our data to contain equal numbers of male and female cells across all genotypes and timepoints. Then, for cells of each sex, read counts were log-normalized and the top 2,000 varying features were identified and analyzed using the FindVariableFeatures function of Seurat. Next, features were selected for downstream integration using the SelectIntegrationFeatures function. Normalized read counts for these features were scaled before performing principal component analysis (PCA). Following this, anchoring features between the two sexes were calculated using FindIntegrationAnchors functions with parameter ‘reduction = “rpca”’ which used reciprocal PCA (top 30 PCs) to identify an effective space in which to find anchors. Finally, samples of the two sexes were integrated with the IntegrateData function using the identified anchoring features.

After integration, expression of these genes was scaled to have a mean at 0 and standard deviation of 1 using the ScaleData function. PCs were analyzed for the expression of these genes using the RunPCA function. The first 30 components were then included for constructing a shared nearest-neighbor graph using FindNeighbors prior to Louvain clustering with FindClusters (resolution set to 0.8). Finally, UMAP dimension reduction was performed using the RunUMAP function with default parameters and 30 dimensions.

To map cells from other genotypes and timepoints to clusters and UMAP space calculated for VPKO Late cells using the methods described above, we used the ‘FindTransferAnchors’ and ‘MapQuery’ functions in Seurat. Transfer anchors between other cells and VPKO Late cells were identified using the ‘FindTransferAnchors’ function with the reference PCA reduction and the first 30 principal components. These anchors were then used to project other cells into the reference embedding with MapQuery, applying the pre-computed reference UMAP model and transferring VPKO Late cluster identities set at a resolution of 0.8.

### Cell Type Assignment

Cell types were identified based on the relative expression of a defined set of renal cell-type specific markers reported in literature and used previously (18,20). The collective expression of each set of cell-type specific marker genes was then scored in each cell using the ‘AddModuleScore_UCell’ function from the UCell package (v2.8.0) (58), with the ‘maxRank’ parameter set to 1500. This scoring method was used as it allows scores for different sets of genes to be compared within the same cell (58). Cell type was assigned for each individual cell by identifying the set of cell-type specific marker genes that attained the highest expression score in that cell.

### PT Class Assignment

Sets of genes exhibiting anti-correlated expression within PT S1, S2, and S3 cells, termed ‘PT Module A’ and ‘PT Module B’, have been described previously (18,20). PT cells were scored for the expression of these sets of genes using the ‘AddModuleScore’ function (59) in Seurat with 100 bins and 50 controls per bin. This scoring method was used as it allows for the expression of the same set of genes to be compared across cells in a dataset. PT Class A cells were identified as those with PT Module A expression score > 0.125 and the others were identified as PT Class B cells.

### Differential Expression (DE) Analysis

To assess the extent of gene regulation caused by *Vhl* and/or *Pbrm1* inactivation, or by time, we performed differential expression analysis using a ‘pseudo-bulking’ approach that treats each mouse, and not each cell, as a biological replicate and thereby reduces the extent of false discoveries in scRNA-seq analysis (60,61). Raw gene expression counts in cells from each mouse were first summed using Seurat’s ‘AggregateExpression’ function to generate a ‘sample × gene’ matrix from a ‘cell × gene’ matrix. The summed data was then used to calculate differential expression using the ‘DESeqDataSetFromMatrix’ function from the DESeq2 package (v1.44.0) (62), using the design formula “∼ Sex + Condition” (where Condition denotes genotype and timepoint) to account for sex-dependent differences in baseline gene expression. Low-count genes were removed prior to analysis, and differential expression was assessed using DESeq2 with Benjamini–Hochberg-adjusted p values. Genes were considered differentially expressed if they were significantly altered (adjusted p value < 0.01) and exhibited a robust effect size (|log2 fold-change| > 0.5).

The following comparisons were made: VKO Early versus ConKO Early, VPKO Early versus VKO Early, VKO Late versus ConKO Late, VPKO Late versus VKO Late, VPKO Late versus ConKO Late, VKO Late versus VKO Early, and VPKO Late versus VPKO Early.

To identify genes that were differentially expressed in cells of one Louvain cluster compared to the others, we used Seurat’s ‘FindMarkers’ function, which performs a Wilcoxon test across all detected genes and corrects for multiple testing using the Benjamini-Hochberg method. The proportion of cells in a cluster in which each gene is detected was also obtained in this analysis.

### Gene Set Enrichment Analysis (GSEA)

GSEA analyses and plotting were performed with the ‘fgsea’ and ‘PlotEnrichment’ functions of the fgsea (63) package (v1.30.0) in R, using gene sets and ranked lists of genes as follows:

A curated list of hypoxia-inducible genes was obtained from the human ‘Hallmark’ gene sets from MSigDB (64). A list of canonical HIFα target genes, validated across multiple human cancer cell lines based on HIFα genome binding (ChIP-seq) and hypoxic regulation (RNA-seq) were obtained from literature (25). Gene names were converted using the ‘orthologs’ function of the babelgene (65) package (v22.9) in R. Genes were ranked in descending order of L2FC in VKO Early versus ConKO Early or VPKO Early versus VKO Early cells.

### Scoring Gene Set Expression

The collective expression of defined sets of genes was evaluated in cells using the ‘AddModuleScore’ function (59) of Seurat, with 100 bins and 50 controls per bin.

For each PT identity, lists of genes that were regulated early (‘Early Up’ and ‘Early Down’) or over time (‘Adaptive Up’ and ‘Adaptive Down’) specifically following *Vhl* inactivation, and whose *Vhl*-dependent regulation was specifically dependent on HIF1A (‘HIF1A Up’ and ‘HIF1A Down’) or on HIF2A (‘HIF2A Up’ and ‘HIF2A Down’), were obtained from literature (20). Cells of each PT identity were evaluated separately for the expression of *Vhl*-dependent or HIFα-isoform specific gene pertinent to their PT identity. Additionally, cells were also scored for the expression of these gene sets irrespective of their PT identity. In this case, the gene sets obtained from the above literature were aggregated to contain every gene that was included as being *Vhl*-dependent or HIFα-isoform specific, in cells of any PT identity.

A list of genes that constitute the integrated stress response (‘ISR Module’) was obtained from literature (30).

### Over-representation Analysis

Over-representation of biological processes GO terms within differentially expressed genes following *Pbrm1* inactivation in *Vhl*-null cells was evaluated using the enrichGO function of the clusterProfiler package (v4.12.6) (66), with multiple testing corrected using false discovery rate. GO terms with recommended restrictions over their use for direct gene product annotation, and those with fewer than 50 genes and more than 1000 genes were excluded from analysis. GO terms were considered to be enriched if they were significantly over-represented (adjusted p < 0.05), and if the average L2FC across all genes that are members of the GO terms was above 0.

### Statistics and Reproducibility

All experiments were performed with at least three biological replicates (mice) per experimental condition. Mice of both sexes were included in every experiment. Where possible, mice from the same litter were used for both ‘early’ and ‘late’ experimental cohorts.

Experiments were repeated in multiple mouse cohorts generated over several years to enhance reproducibility of the findings. No statistical method was used to predetermine sample size. No data were excluded from the analyses. The experiments were not randomized. Except for manual counting of tdTomato-positive and BrdU-positive cells by IHC, the investigators were not blinded to allocation during experiments and outcome assessment.

For differential expression testing between genotypes and timepoints, p values were calculated using Wald tests performed in the ‘results’ function of DESeq2 in R using an absolute-effect alternative hypothesis (altHypothesis = “greaterAbs”; lfcThreshold = 0; independentFiltering = FALSE), and adjusted p values were evaluated at ‘alpha = 0.01’. The p value was corrected for multiple testing using the Benjamini-Hochberg method. For DE testing between Louvain clusters, p values were calculated using the Wilcoxon test as part of the ‘FindMarkers’ function in Seurat, with multiple testing corrected by the Benjamini-Hochberg method.

For over-representation analysis, p values were calculated using the hypergeometric test as part of the enrichGO function in the clusterProfiler package in R.

Gene set expression scores were compared across genotypes and timepoints using either a one-way or two-way two-tailed ANOVA test in R, with multiple testing correction using Benjamini-Hochberg method.

For statistical analyses of cluster composition, cluster proportions were calculated on a per-mouse basis, treating each mouse as an independent biological replicate. Per-mouse proportions were compared between conditions using pairwise Wilcoxon rank-sum tests in R with p values adjusted for multiple testing using the Benjamini–Hochberg method.

IHC and IF quantification data were analyzed between genotypes and timepoints using two-tailed two-way ANOVA in R. p values were corrected for multiple testing using the Benjamini-Hochberg method. Differences in frequency distributions were tested using multivariate analysis of variance with the ‘manova’ function in R and the Wilks’ statistic. Resulting *p* values were adjusted for multiple testing using the Benjamini-Hochberg method.

To quantify whether transcriptional changes in VPKO Late cells could be explained by additivity of *Vhl* and *Pbrm1* effects, the observed differential expression in VPKO Late (VPKO Late versus ConKO Late) was compared to an expected change predicted from the sum of two independently estimated components: the effect of *Vhl* inactivation at late timepoint (VKO Late versus ConKO Late) and the effect of *Pbrm1* inactivation measured early (VPKO Early versus VKO Early). Deviations from additivity were quantified as a residual (Residual = Total − Predicted), with uncertainty propagated from the standard errors (SE) of the component changes – SE_residual = √(SE_total² + SE_Vhl² + SE_Pbrm1²), where SE_total is from VPKO Late vs ConKO Late, SE_Vhl from VKO Late vs ConKO Late, and SE_Pbrm1 from VPKO Early vs VKO Early. Residuals were converted to z-statistics (z = Residual/SE_residual) and assessed using two-sided p values. Analyses were focused on genes robustly and significantly altered in at least one of the component or total contrasts, as defined by the thresholds used for pseudo-bulked differential expression.

Spearman coefficients for correlation (ρ) between differential gene expression in different genotypes using the ‘cor.test’ function in R. Only genes that were significantly and robustly regulated in either of the differential expression analyses being compared were included in the analysis.

### Data Visualization

Single-cell sequencing data was visualized with the in-built functions of pheatmap (v1.0.12) (67), and ggplot2 (v3.5.1) (68) packages in R (v4.3.2 and v4.4.1) (69).

### Data Availability

The scRNA-seq data for *Vhl*-null cells (ConKO and VKO) harvested early and late after recombination data used in this study are available in the Gene Expression Omnibus (GEO) database under accession code GSE253168 [https://www.ncbi.nlm.nih.gov/geo/query/acc.cgi?acc=GSE253168]. The scRNA-seq data for VPKO and ConPKO mice harvested early or late after recombination are available in the Gene Expression Omnibus (GEO) database under accession code GSE319776 [https://www.ncbi.nlm.nih.gov/geo/query/acc.cgi?acc=GSE319776].

## Acknowledgements

Funding for the work was received from the Oxford Branch of the Ludwig Institute for Cancer Research. This work was also supported by the Francis Crick Institute, which receives its core funding from Cancer Research UK (FC001501), the UK Medical Research Council (FC001501), and the Wellcome Trust (FC001501). S.K was sponsored by a Christopher Welch Scholarship and the Clarendon Fund. We thank Norma Masson (Nuffield Department of Medicine, Oxford) for performing the tdTomato and PBRM1 immunoblotting, Indrika Ratnayaka (Ludwig Institute for Cancer Research, Oxford) for helping with cryostat sections, Ran Li (Nuffield Department of Medicine, Oxford) for bioinformatics support, and Ruddy Montandon, Ghada Ben Youssef and Mohammed Islam (Center for Human Genetics, Oxford) for flow cytometry services. We also thank all staff in the Functional Genetics Unit, Center for Human Genetics, Oxford for their support with animal husbandry.

## Author Contributions

S.K, J.D.C.C.L, and J.A performed mouse husbandry, tamoxifen and BrdU administration and harvests. S.K, J.D.C.C.L, and N.K processed tissues and performed tdTomato IHC. J.D.C.C.L performed HE and Oil Red O staining, and PBRM1, HIF1A and HIF2A IHC. S.K and J.D.C.C.L performed dual IHC, tdTomato/PAS staining, and tdTomato/BrdU/PAS staining. S.K performed IF. S.K performed quantitative analysis of IHC data. S.K and J.D.C.C.L prepared scRNA-seq samples and libraries. S.K and D.R.M processed scRNA-seq data and designed analyses. S.K analyzed scRNA-seq data. C.W.P, D.R.M, and J.A provided expert insights and guidance to the project. P.J.R conceptualized the study and obtained funding. S.K, D.R.M, J.A, and P.J.R supervised the project. S.K, J.D.C.C.L, and P.J.R wrote the manuscript. C.W.P, D.R.M, and J.A reviewed and edited the manuscript.

