## Supplementary_Legends for "Discrete genetic effects of *VHL* and *PBRM1* inactivation co-operate to disrupt epithelial homeostasis and promote ccRCC"

### Supplementary Figure Legends

**Supplementary Fig. 1: Co-deletion of *Pbrm1* in an oncogenic 'cell tagging' model of *Vhl* inactivation.**

**(a)** Schematic diagram depicting the design of the $Vhl^{pjr.fl}$ and $Pbrm1^{fl}$ alleles, and genotypes produced following tamoxifen (TMX) inducible Cre-mediated recombination of the two alleles. In addition to $Vhl^{pjr.fl}$, ConPKO mice carry a wild-type $Vhl^{wt}$ allele, while VPKO mice carry a constitutively inactivated and untagged $Vhl^{jae.KO}$ allele. P, Promoter; U, untranslated region; E, Exon; I, Intron; pA, polyadenylation site; P2A, porcine teschovirus 2A peptide; SA, splice acceptor.

**(b)** Representative density plots depicting the gating strategy to isolate live, single, tdTomato-positive cells from dissociated kidney cell suspensions. Cells of interest were distinguished from debris by gating on Forward Scatter Area (FSC-A) versus *S*ide Scatter area (SSC-A). Single cells were distinguished from doublets and aggregates by gating the cells of interest on FSC-A versus Forward Scatter Width (FSC-W). Live cells were identified as single cells not stained positive by DAPI, which was detected using a 405-450_50 band/pass filter. tdTomato-positive cells were isolated from the live, single cell population based on the positive signal detected with the 561-582_15 band/pass filter.

**(c)** Genomic PCR for *Pbrm1* alleles performed on FAC-sorted tdTomato-positive cells from VKO and VPKO mice given 5x 2 mg tamoxifen and harvested early (1-3 weeks) after recombination. Amplicon sizes for $Pbrm1^{wt}$ (243 bp) and $Pbrm1^{fl}$ (400 bp) are indicated.

**(d)** Representative immunoblots (IB) for tdTomato protein or PBRM1 in FAC-sorted tdTomato-negative (−) or tdTomato-positive (+) cells from VKO and VPKO mice given 5 × 2 mg tamoxifen and harvested early (1-3 weeks) following recombination.

**(e)** Representative H&E staining (left) and tdTomato IHC (right) performed on serial sections of renal tumors in VPKO mice (n = 4) observed at 12-17 months following recombination (aged timepoint). Insets depict cytoplasmic clearing in the tumors (left, upper) compared to strong eosin staining of a morphologically normal tubule (left, lower) as well as uniform tdTomato-positive staining of cells within tumors (right). Scale bars, 100 µm. 40x magnification.

**(f)** Representative IHC for HIF1A, HIF2A and PBRM1 renal tumors in VPKO mice (n = 4) observed at 12-17 months following recombination (aged timepoint) demonstrating HIFα stabilization and loss of PBRM1. Scale bars, 100 µm. 40x magnification.

**Supplementary Fig. 2: Effect of *Pbrm1* inactivation on *Vhl* and HIFα-dependent gene expression in PT cells.**

**(a)** Expression of sets of genes up- or downregulated in cells of different PT identities in a HIF1A- ('HIF1A Up' or 'HIF1A Down') or HIF2A-dependent manner ('HIF2A Up' or 'HIF2A Down') in ConKO, VKO, and VPKO cells of different PT identities sequenced early (1-3 weeks) after recombination. The PT identities of scored cells are denoted by labels on the right side of the plot.

**(b)** Expression of sets of genes up- or downregulated in cells of different PT identities early after *Vhl* inactivation ('Early Up' or 'Early Down') in ConKO, VKO, and VPKO cells of different PT identities sequenced 1-3 weeks (early) following recombination.

**(a, b)** Median values and inter-quartile ranges of the data are presented separately for cells from each mouse. Pairwise comparisons have been made between means of the median values for mice of different genotypes using two-tailed one-way ANOVA with multiple testing correction using the Benjamini-Hochberg method. For cells of PT identities in which specific HIF1A-dependent, HIF2A-depedenent or early regulated *Vhl*-dependent genes had not been identified, the module score has been assigned a value of 0.

**Supplementary Fig. 3: Sustained and independent transcriptional effects of *Pbrm1* inactivation in PT cells.**
**(a)** UMAP plot depicting ConKO, ConPKO, VKO, and VPKO cells harvested early (1-3 weeks) following recombination, showing that ConPKO cells occupy distinct UMAP space to ConKO, VKO, or VPKO cells.
**(b)** Cell clusters occupied by ConKO, ConPKO, VKO, and VPKO cells in UMAP space (left panel) and the proportion of cells from mice of each genotype that are present in each cluster (right panel). Data are presented as median values, with the inter-quartile range indicated by error bars.

**Supplementary Fig. 4: Emergent cell states among *Vhl/Pbrm1*-null cells.**
**(a)** UMAP plots depicting PT type (left) and class (right) of VPKO Late cells, showing that the distribution of VPKO Late cells in UMAP space is largely associated with their PT type and class.
**(b-d)** Pseudo-bulked L2FC plotted against the negative logarithm of the p value (two-tailed Wald test with multiple testing correction by Benjamini-Hochberg method) for genes in Cluster 12 versus all other clusters combined **(b)**, Cluster 9 versus all other clusters combined **(c)**, and Cluster 2 versus all other clusters combined (**d**). Genes are colored by whether they exhibited 'emergent' regulation in VPKO Late cells, as analyzed in Fig. 3a.

**Supplementary Fig. 5: Maintenance of a *Vhl*-dependent proliferative drive following *Pbrm1* inactivation.**
**(a, b)** Representative tdTomato IHC counterstained with hematoxylin in the renal papilla **(a)** or renal cortex **(b)** of ConKO, VKO, ConPKO and VPKO mice given 5x 2 mg tamoxifen and harvested early (1-3 weeks) or late (4-12 months) following recombination. Scale bars, 100 µm; 40x magnification.
**(c)** Representative immunofluorescence staining of tdTomato (yellow) and Ki67 (magenta), counterstained with DAPI (cyan) in the renal cortex, as used for automated quantification. Examples of tdTomato-positive, Ki67-positive dual-labelled cells are denoted by white arrows Scale bars, 50 µm. 20x magnification.
**(d)** Automated quantification (see '**Methods**') of the proportion of tdTomato-positive cells that are also Ki67-positive in the renal cortex of ConKO, ConPKO, VKO, and VPKO mice harvested early (1-3 weeks) and late (4-12 months) following recombination. Error bars indicate Mean ± SE. Pairwise comparisons were made by two-tailed, two-way ANOVA test with multiple testing correction using the Benjamini-Hochberg method. The number of biological replicates (mice) analyzed for each genotype and timepoint is indicated.

**Supplementary Fig. 6: Morphological alterations in *Vhl/Pbrm1*-null cells.**

**(a, b)** Representative images of dual IHC for tdTomato (brown) and BrdU (grey) coupled to PAS staining (pink) counterstained with hematoxylin in kidneys of ConPKO mice (n = 5) harvested late (4-12 months) following recombination (**a**), and in VKO mice (n = 5) harvested early (1-3 weeks) after recombination (**b**). Luminal, crowded, and multi-layered tdTomato-positive cells that have incorporated BrdU are present in kidneys from ConPKO Late mice and are denoted by dashed black arrows. tdTomato-positive cells that have incorporated BrdU exhibit normal morphology and are present within the tubular boundary in VKO Early mice. Scale bars, 50 µm. 40x magnification.

#### Supplementary Table Legends

**Supplementary Table 1:** Pseudo-bulked differential gene expression in PT cells following *Vhl* and/or *Pbrm1* inactivation early after recombination, late after recombination, or over time.

**Supplementary Table 2:** Differential gene expression in Clusters 12, 9, or 2 compared to all other clusters combined, as defined for VPKO Late cells and to which cells from other genotypes and timepoints are projected.
