## Supplementary figures and images for "Discrete genetic effects of *VHL* and *PBRM1* inactivation co-operate to disrupt epithelial homeostasis and promote ccRCC"

### Supplementary_Figure_1

**Supplementary Fig. 1**

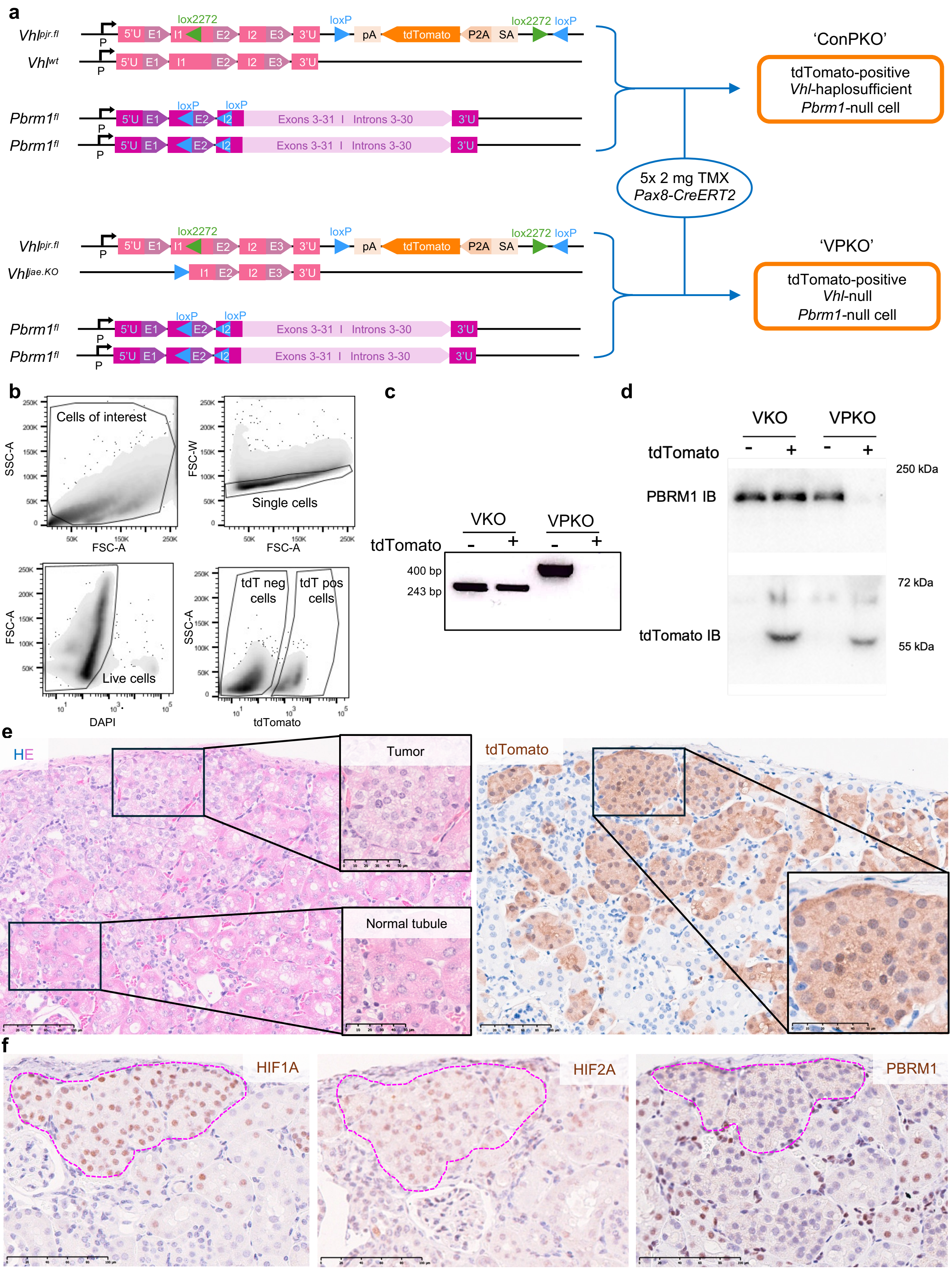

### Supplementary_Figure_2

**Supplementary Fig. 2**

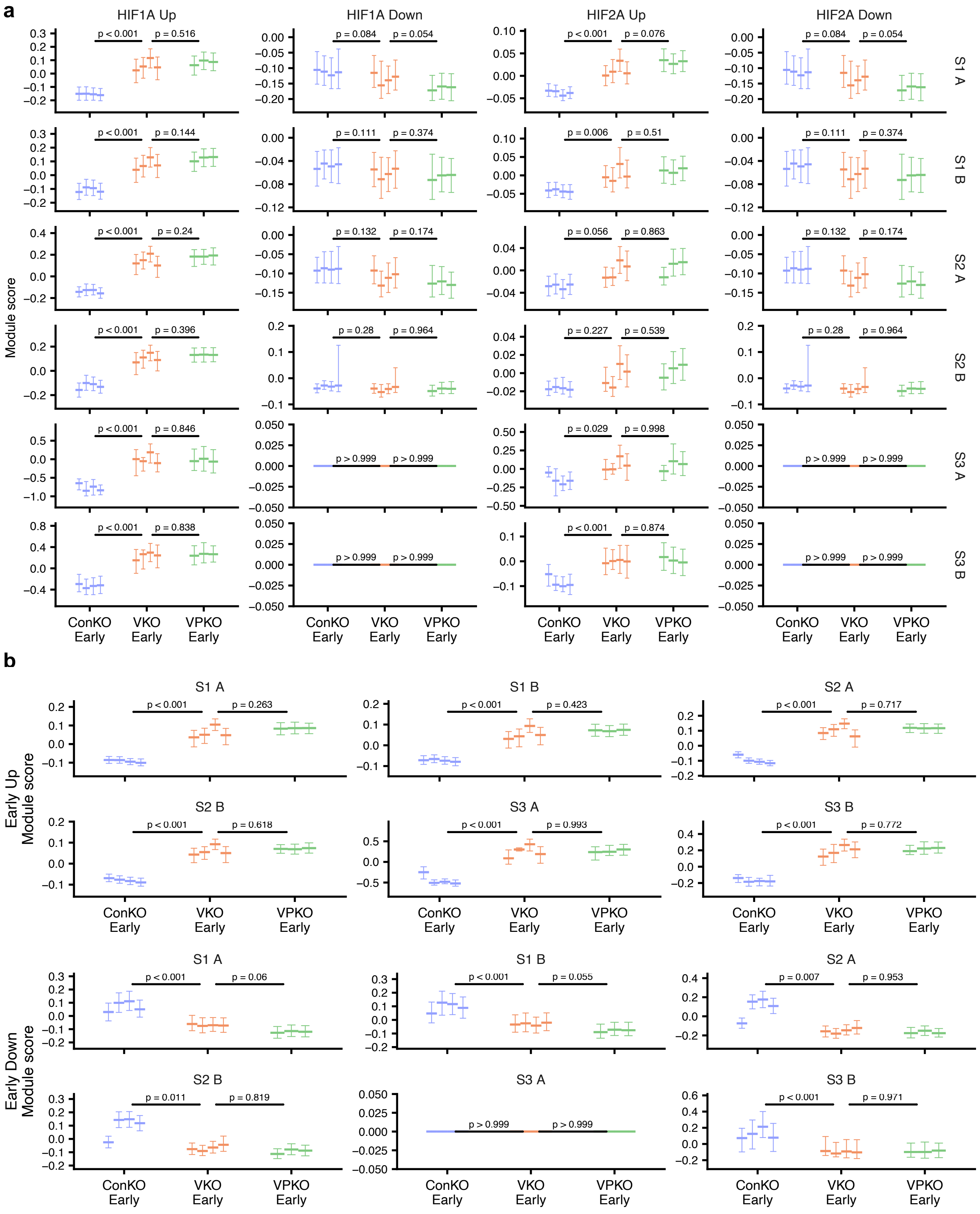

### Supplementary_Figure_3

#### Supplementary Fig. 3

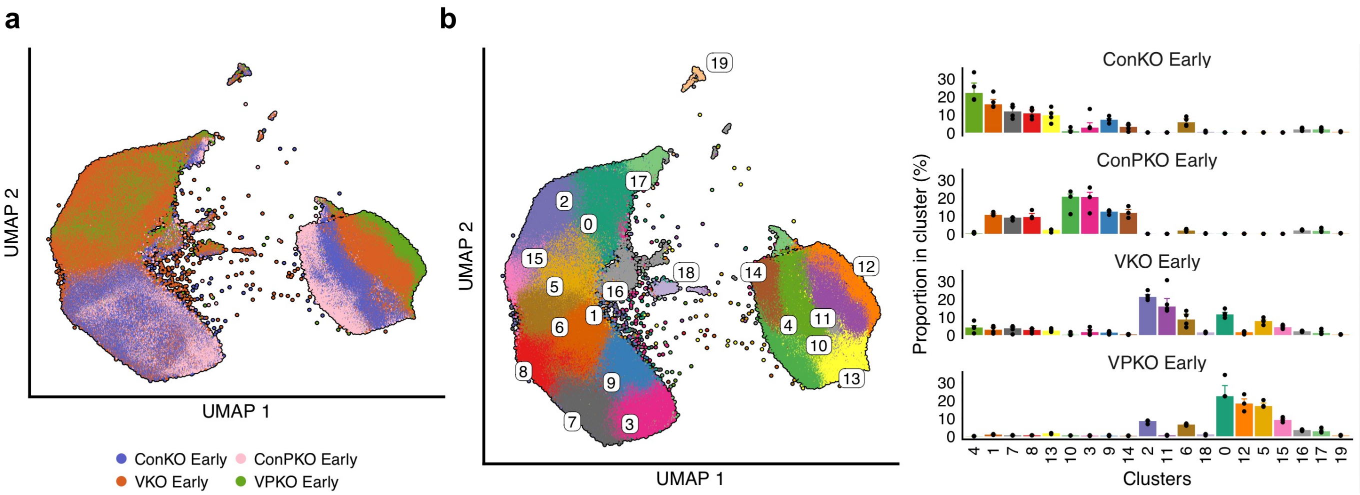

### Supplementary_Figure_4

#### Supplementary Fig. 4

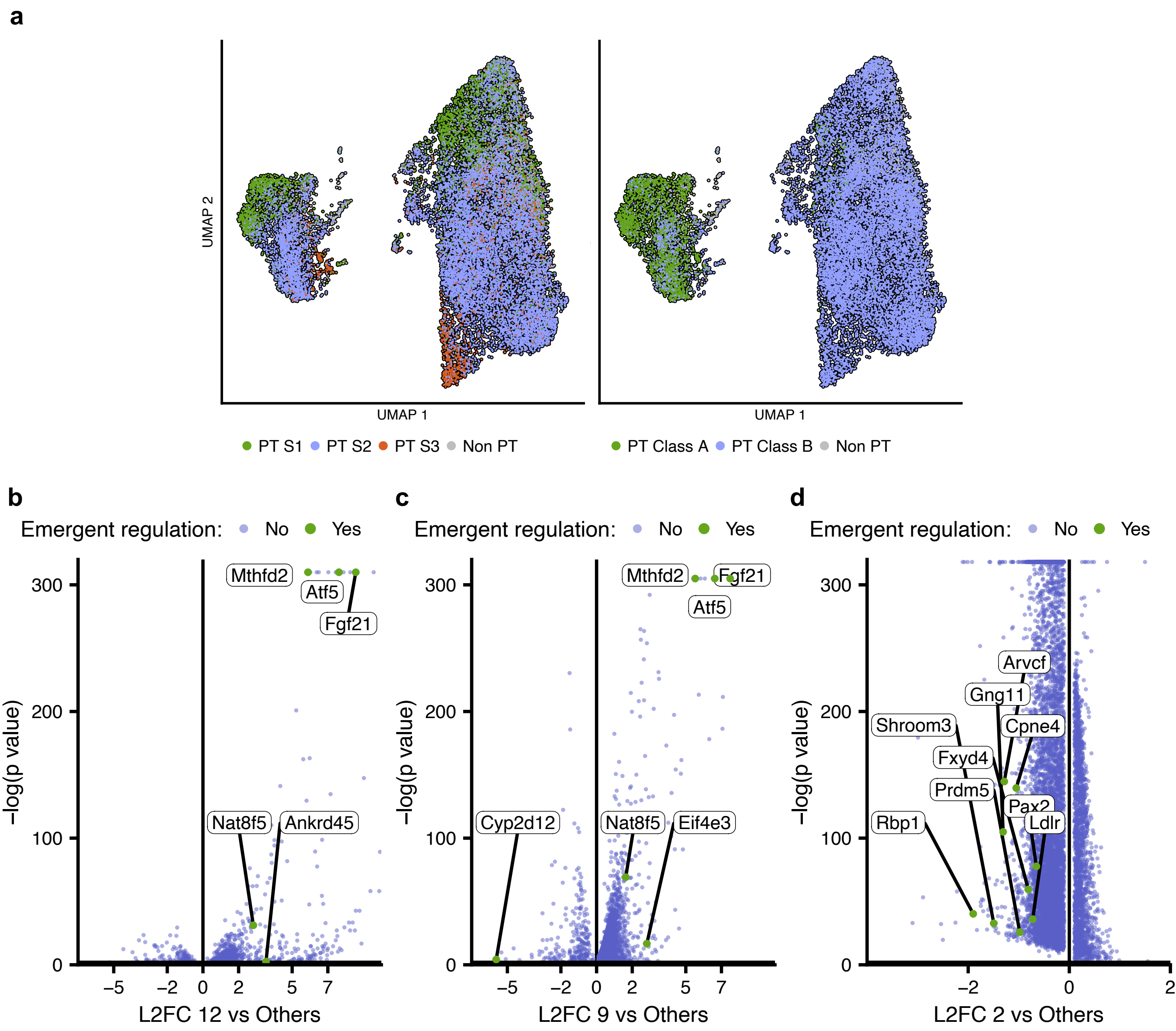

### Supplementary_Figure_5

#### Supplementary Fig. 5

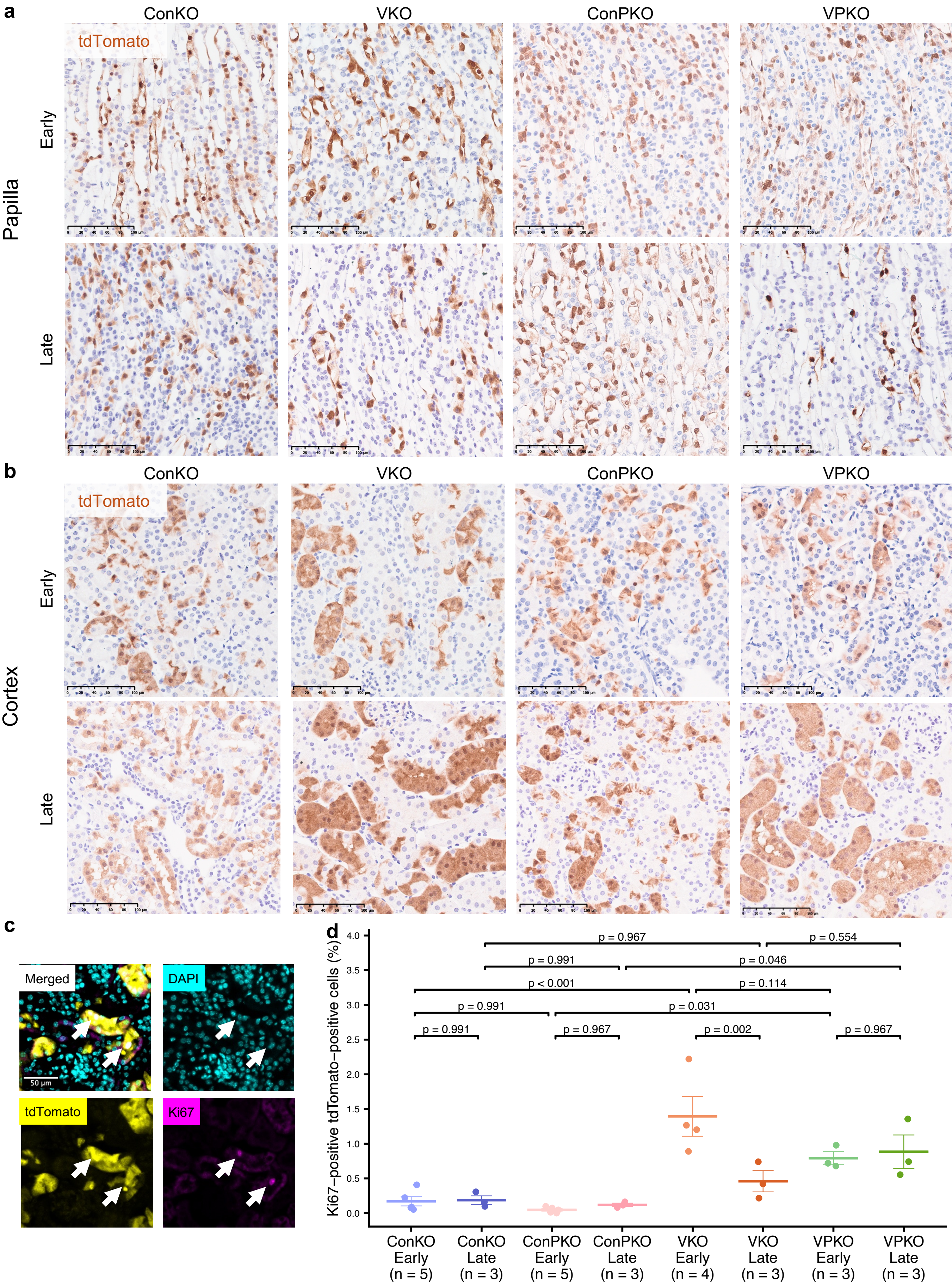

### Supplementary_Figure_6

##### Supplementary Fig. 6

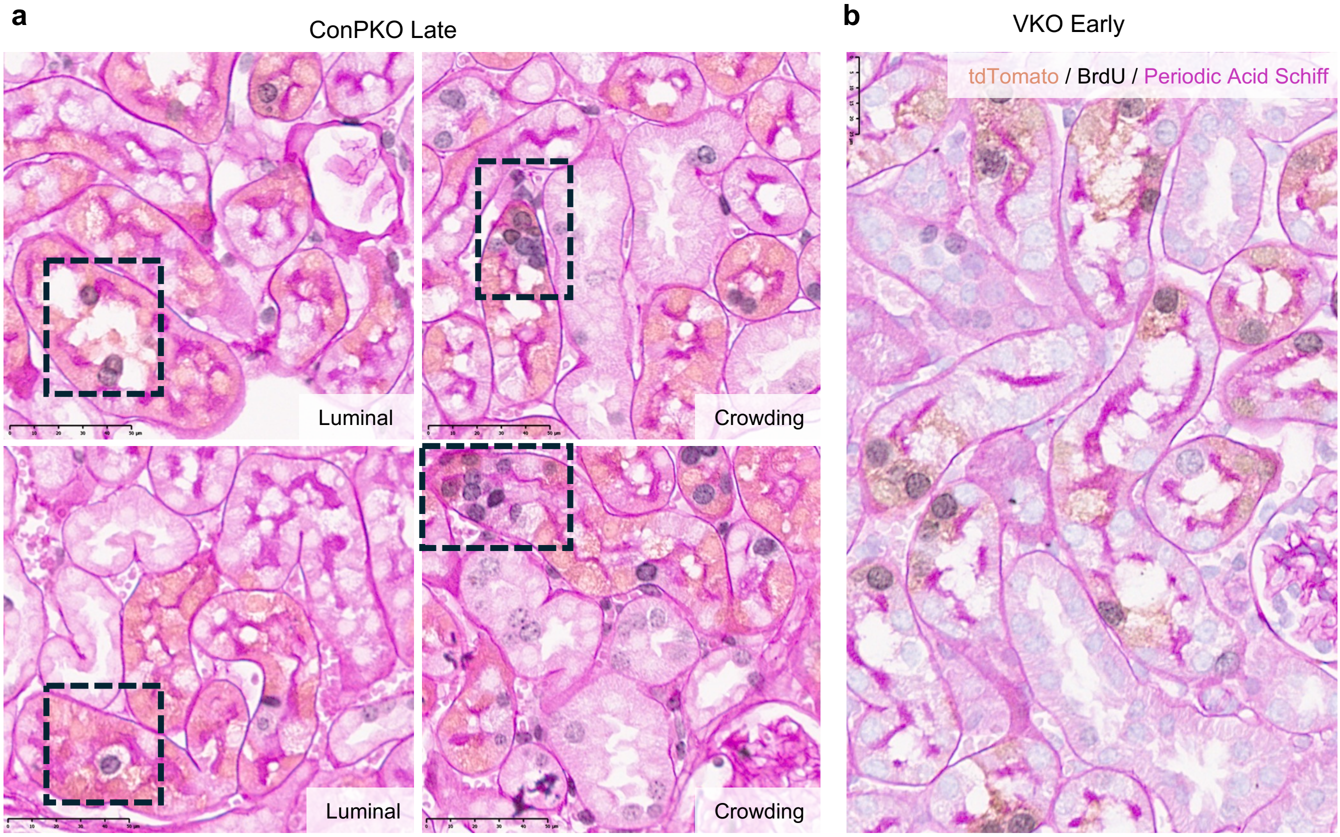
